# Neural Spectral Prediction for Structure Elucidation with Tandem Mass Spectrometry

**DOI:** 10.1101/2025.05.28.656653

**Authors:** Runzhong Wang, Mrunali Manjrekar, Babak Mahjour, Julian Avila-Pacheco, Joules Provenzano, Erin Reynolds, Magdalena Lederbauer, Eivgeni Mashin, Samuel Goldman, Mingxun Wang, Jing-Ke Weng, Desirée L. Plata, Clary B. Clish, Connor W. Coley

## Abstract

Structural elucidation using untargeted tandem mass spectrometry (MS/MS) has played a critical role in advancing scientific discovery [1, 2]. However, differentiating molecular fragmentation patterns between isobaric structures remains a prominent challenge in metabolomics [3–10], drug discovery [11–13], and reaction screening [14–17], presenting a significant barrier to the cost-effective and rapid identification of unknown molecular structures. Here, we present a geometric deep learning model, ICEBERG, that simulates high-energy collision-induced dissociation in mass spectrometry to generate chemically plausible fragments and their relative intensities with awareness of collision energies and polarities. We utilize ICEBERG predictions to facilitate structure elucidation by ranking a set of candidate structures based on the similarity between their predicted *in silico* MS/MS spectra and an experimental MS/MS spectrum of interest. This integrated elucidation pipeline enables state-of-the-art performance in compound annotation, with 40% top-1 accuracy on the NIST’20 [M+H]^+^ adduct subset and with 92% of correct structures appearing in the top ten predictions in the same dataset. It achieves 46% top-1 and 86% top-10 accuracies when tested on the open-access MassSpecGym benchmark, and outperforms SIRIUS on a recently released test set with previously uncharacterized structures. We demonstrate several real-world case studies, including identifying clinical biomarkers of depression and tuberculous meningitis, annotating an aqueous abiotic degradation product of the pesticide thiophanate methyl, disambiguating isobaric products in pooled reaction screening, and annotating biosynthetic pathways in *Withania somnifera*. Overall, this deep learning-based paradigm for structural elucidation enables rapid molecular annotation from complex mixtures, driving discoveries across diverse scientific domains.

## Introduction

From metabolomics [1–10] to drug discovery [11–13] and reaction screening [14–17], the identification of molecular structures using liquid chromatography with tandem mass spectrometry (LC-MS/MS) has broad applications in chemistry and biology. In a structural elucidation campaign with LC-MS/MS, the sample mixture is first separated using liquid chromatography. At each retention time (RT), compounds eluting from the LC column are softly ionized and separated by different mass-over-charge (*m/z*) values, i.e., separated by MS1. Each precursor ion within a certain RT and *m/z* window can then be fed into a second fragmentation stage where the bonds are cleaved and the fragments are detected as peaks, cumulatively called a tandem mass spectrum (MS/MS) [18]. Different analytical workflows may employ different fragmentation mechanisms, ionization modes, and mass detectors; here, we focus on high-energy collision-induced dissociation (HCD) using Orbitrap mass spectrometers. In the broader context of analytical chemistry, mass spectrometry plays a unique role due to its exceptional sensitivity, precision, and throughput [19].

The ultimate goal of structure elucidation directly from MS/MS spectra remains a formidable challenge [20]. Current practices overwhelmingly rely on matching experimental MS/MS spectra to MS/MS reference spectral libraries of known structures. However, these reference libraries either require authentic standards or laborious orthogonal structure elucidation—both of which exist only for a vanishingly small fraction of the vast space of all possible molecules. In real-world metabolomics studies, most MS/MS features measured do not have any close match to known standards, known as “metabolite dark matter” [21, 22].

We tackle structural elucidation of such dark matter by building upon the recent success of heuristic [23–26] and machine learning-based [27–30] approaches that predict MS/MS from molecular structures. These prediction approaches mitigate the challenge of limited reference library sizes by generating *in silico* spectra for candidate structures that may not exist in these libraries. These predicted MS/MS spectra can be used to narrow down the candidate structures for an unannotated spectrum, reducing the burden of acquiring standards for experimental validation of a hypothesized structural annotation (Fig. 1a). The ability to predict MS/MS spectra directly from molecular structures illuminates new areas of chemical space that were previously limited by spectral libraries. For example, structure libraries (e.g., HMDB [31], PubChem [32]), enumerative techniques (e.g., virtual reaction enumeration [33]), and generative modeling (e.g., mimicking the distribution of known metabolites [34]) could provide plausible candidate structures. Such a structural elucidation workflow leveraging MS/MS prediction can significantly accelerate the analysis of large numbers of unannotated spectra, opening up new frontiers in mass spectrometry-based discoveries. An alternative strategy for retrieval is predicting molecular fingerprints from MS/MS spectra [35–37], but we show that the leading method, SIRIUS [37], empirically underperforms our *in silico* fragmentation models. We acknowledge that there also exist complementary methods that focus on *de novo* structural prediction from MS/MS without the use of an explicit MS/MS predictor [38–42], which are as yet limited in terms of accuracy [43]. We are also aware of the challenge of in-source fragmentation leading to false positive features [44], which should be detected and mitigated by experimental setting and retention time correlation [45], therefore beyond the scope of this paper that focuses on the downstream structural elucidation task.

**Fig. 1:**
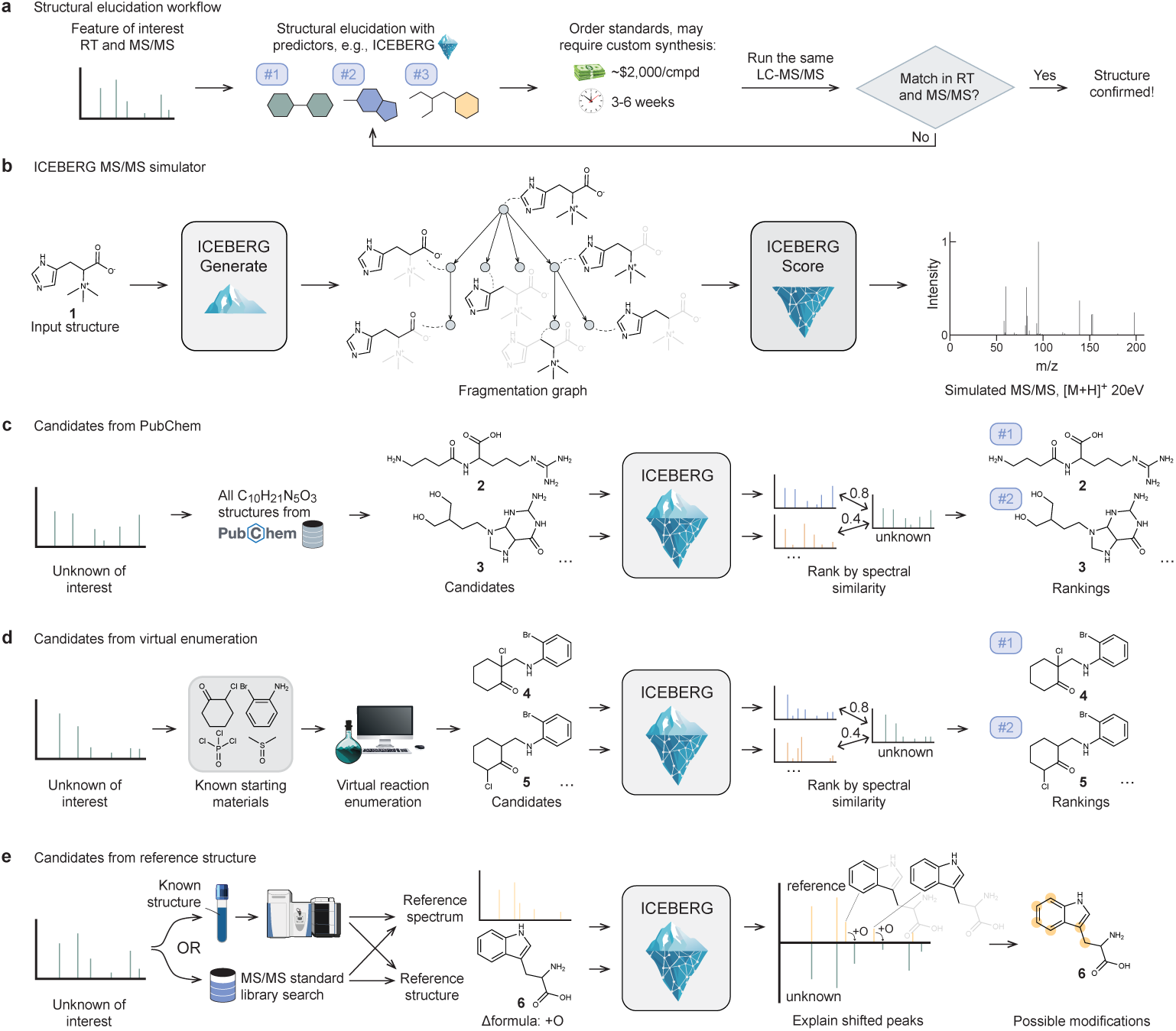
ICEBERG is an *in silico* predictor of tandem mass spectrometry and accelerates structural elucidation with different sources of candidates. a, Structural elucidation from MS/MS is a time-intensive and costly task, especially when custom synthesis of authentic standards is needed, as shown in this prototypical structural elucidation workflow. **b**, ICEBERG is a two-stage neural network model that simulates the fragmentation process in LC-MS/MS. ICEBERG-Generate first generates a fragmentation graph given a molecular structure. ICEBERG-Score then predicts the intensities for each fragment, together with other inputs such as adduct type and collision energy. Structural elucidation is achieved by using ICEBERG-generated MS/MS to rank candidate structures, where candidates can be defined through several complementary strategies; three are demonstrated as follows. **c**, If the unknown is expected to be in a chemical library, e.g., PubChem, and the chemical formula is known or can be inferred, all structures in PubChem with the same chemical formula could be treated as candidates. **d**, If the unknown structure is novel, there might be prior knowledge that helps constrain the set of possible structures, for example, through virtual reaction enumeration given known starting materials. **e**, If the unknown structure is thought or found to be closely related to a reference structure (e.g., through late-stage functionalization from chemical or biological sources), where the MS/MS of the unknown is close to the reference’s, candidates could be derived from the reference structure. For example, with a known formula difference of +O, ICEBERG identifies substructures that explain peaks shifted by +O and highlights the most plausible sites of hydroxylation.

Inferring Collision-induced-dissociation by Estimating Breakage Events and Reconstructing their Graphs (ICEBERG) is a geometric deep learning model that simulates mass spectra from structures that we previously introduced in Goldman et al. [46]. The model design is inspired by the physical fragmentation process in tandem mass spectrometers. ICEBERG is a two-stage model where the first model (ICEBERG-Generate) generates the most plausible molecular fragmentations, creating a fragmentation graph in an auto-regressive manner; given the highest probability fragments, the second model, ICEBERG-Score, predicts intensities and hydrogen shifts for each fragment (Fig. 1b), yielding the final MS/MS prediction. With predictions from both stages, the model offers physical grounding on the fragment structures together with the intensities.

In this work, we introduce a substantially improved version of ICEBERG as a state-of-the-art neural MS/MS predictor for small organic molecules (Extended Data Fig. 1). We expand the relevant application space by including collision energy and ionization polarity (positive and negative mode) as key parameters. Further, we introduce new innovations in model design, such as contrastive fine-tuning [47], accounting for possible charge migrations in fragmentation [48], and utilizing spectral entropy [49] in training and retrieval. These technical improvements together contribute to an increase in top-1 annotation accuracy from 25% to 40% on the NIST’20 benchmark, while also surpassing the off-the-shelf structural elucidation software, SIRIUS [37]. To demonstrate how these improvements have impacted elucidation, we showcase successful applications with different sets of candidate structures: ICEBERG helps to identify three novel biomarkers from clinical studies and identify a pesticide degradation product with candidates from PubChem (Fig. 1c), elucidate isobaric products in pooled reaction screening with candidates from virtual reaction enumeration (Fig. 1d), and annotate structures in a biosynthetic pathway with candidates from reference structures (Fig. 1e). Collectively, these case studies illustrate how the ICEBERG approach to MS/MS prediction enables structure elucidation in diverse scientific settings. Finally, the open-source philosophy of ICEBERG may help democratize access for the scientific community in annotating structures from LC-MS/MS.

## Results

### *In silico* validation of MS/MS prediction accuracy through quantitative benchmarking

We evaluate ICEBERG on Orbitrap HCD spectra collected from the NIST’20 dataset, a library developed by the National Institute of Standards and Technology (NIST) [51]. Details of ICEBERG’s architecture and hyperparameters can be found in Methods. Following prior work, we reserve ∼10% of reference spectra for testing and ∼10% for validation. Generalization is assessed through two types of splits. In the random split, structures with different InChI keys are randomly put into training, validation, and testing sets (Fig. 2a). In the scaffold split, structures with the same molecular scaffold are kept in the same set (Extended Data Fig. 2b), providing a more challenging out-of-distribution evaluation that assesses the ability of the model to generalize to distinct structural families. For both splits, we evaluate the spectral prediction accuracy in terms of the average cosine similarity of the predicted and ground truth spectra, as well as retrieval accuracy to assess models’ performance in elucidation given a set of candidate isomers from PubChem. We also report ICEBERG on the open-source MassSpecGym benchmark [43] which emphasizes the generalization ability to dissimilar molecules (Fig. 2b), whose training, testing, and validation structures must have at least an MCES (maximum common edge subgraph) edit distance of 10. Further detailed evaluations can be found in Supplementary Section S2.

**Fig. 2:**
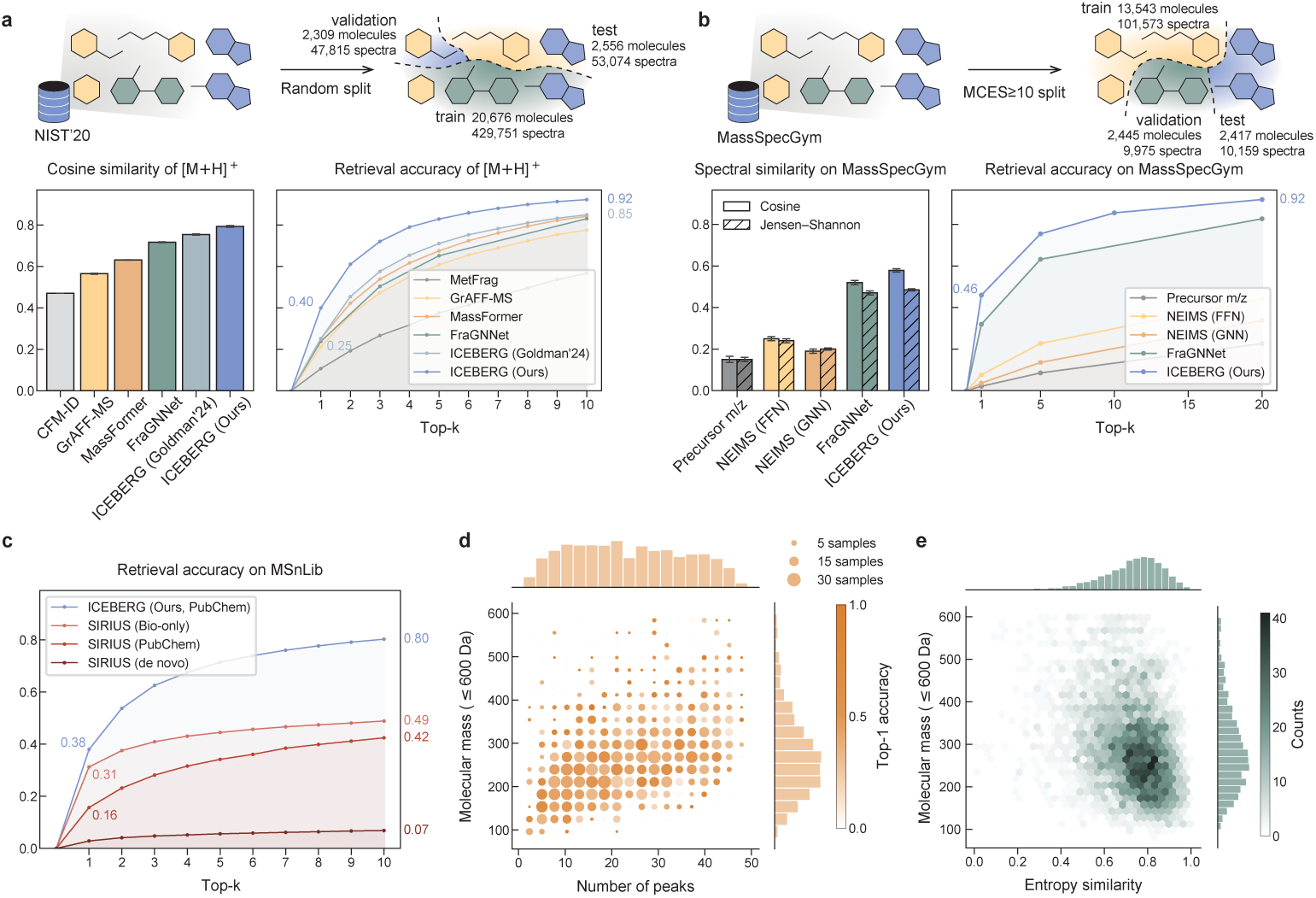
Quantitative evaluation of ICEBERG demonstrates state-of-the-art accuracy in MS/MS prediction and structure annotation. **a**, ICEBERG predicts spectra more accurately on the NIST’20 dataset compared to other prediction tools as measured by cosine similarity as well as structural elucidation through database retrieval. Error bars with 95% confidence interval are reported for 3 random seeds for cosine similarity. **b**, ICEBERG also outperforms peer methods on MassSpecGym dataset [43] with a split that ensures at least an MCES edit distance of 10 between any training, validation, and testing structures, evaluating the generalization ability to unseen molecular scaffolds as a more challenging setting. The term Jensen–Shannon similarity is used in the MassSpecGym benchmark for entropy similarity [49]. Following the benchmark, we report 99.9% confidence intervals upon bootstrapping (20,000 resamples). **c**, ICEBERG outperforms SIRIUS [37] on a subset of MSnLib [50] whose structures have not been reported in NIST or GNPS. We tested the retrieval accuracy of SIRIUS with either PubChem candidates or candidates from bio-relevant libraries, and its accuracy for *de novo* generation. **d**, Retrieval accuracy as a function of the molecular mass and number of peaks in the experimental spectrum shows that ICEBERG performance is consistent with respect to different sizes of molecules and various numbers of peaks. **e**, Spectral similarity as a function of molecular mass and the spectral entropy similarity [49] to an experimental spectrum. The slight negative correlation is attributable to the greater number of peaks larger molecules tend to provide, negatively biasing the entropy similarity metric.

The architectural and training modifications improve ICEBERG by a substantial margin from 25.1% (95% CI 23.5–26.7%) top-1 [M+H]^+^ retrieval accuracy to 40.0% (95% CI 39.2–40.8%) on the random split of NIST’20 (Fig. 2a) and from 31.9% (99.9% CI 30.4–33.5%) to 46.0% (99.9% CI 44.3–47.6%) on the MassSpecGym benchmark (Fig. 2b). Additional quantitative results show performance on the scaffold split and 6 a wider range of adducts for which not all baseline methods are able to make predictions [30] (Extended Data Fig. 2). We benchmark heuristic-based fragmentation methods while excluding comparisons to certain other baseline models that do not permit retraining (Supplementary Section S2.1). On the MassSpecGym benchmark, all previously reported baselines are considered, where “Precursor m/z” is a trivial baseline that simply predicts a single-peak spectrum at the precursor m/z. All technical contributions are validated by an ablation study that confirms the positive contribution of each design choice to overall performance (Extended Data Table 1).

We also demonstrate the superiority of ICEBERG as a structural elucidation tool over the off-the-shelf software, SIRIUS [37] (Fig. 2c). The evaluation is performed on a recently released MS/MS dataset, MSnLib [50]. The testing set contains 8,519 MS2 spectra that have not been reported in NIST or GNPS, ensuring no overlap with the training data of ICEBERG and SIRIUS. ICEBERG takes the full PubChem database as the candidate set (over 100 million structures) and achieves 38.0% top-1 accuracy and 80.3% top-10 accuracy, surpassing SIRIUS, either taking a bio-relevant subset (*<*1 million structures) or the full PubChem set. *De novo* generation in theory covers the entire chemical space, but its accuracy is inferior to retrieval-based methods. The top-10 identification accuracy of ICEBERG is almost doubled compared to SIRIUS, suggesting our robustness and real-world utility. It is worth noting that a perfect top-1 accuracy remains challenging due to certain ambiguous isomers that MS/MS cannot distinguish, e.g., leucine vs isoleucine.

ICEBERG-generated spectra provide consistently accurate predictions across a range of peak counts and molecular masses, achieving robust retrieval accuracies (Fig. 2d) and similarities to experimental spectra (Fig. 2e). We attribute the comparatively strong performance on large structures to the model’s ability to reason about chemical structure through graph neural networks, whereby the geometric deep learning module generalizes across molecular sizes. We observe a difference in accuracy across different chemical classes (Extended Data Fig. 3a). Model performance across several adduct types shows that higher performance is achieved for adducts that are more abundant in the dataset (Extended Data Fig. 3b).

ICEBERG predicts fragmentation spectra in an inherently explanatory manner that attributes each peak to a substructural fragment, yet the veracity of these explanations is not straightforward to assess. Experimental spectra do not typically reveal such detailed structural information about each peak [52]. However, we evaluate the consistency of ICEBERG’s explanations with intuition and with inductive cleavage energy [53]. The trend of model predictions across different collision energies shows that ICEBERG learns to break “weaker” C-O bonds at lower collision energies and learns to break “stronger” aromatic bonds at higher collision energies (Extended Data Fig. 4a). Collision energy-dependent fragmentation further aligns with density functional theory (DFT)-level calculations of cleavage energy of the same fragmentation event (Extended Data Fig. 4b). Qualitatively, *in silico* spectra are predicted to have more fragmentation as collision energy increases (Extended Data Fig. 4c).

### Elucidation of biomarkers in clinical metabolomics studies of depression

We set out to demonstrate how ICEBERG can be used effectively in a structure elucidation pipeline across a variety of small-molecule compound classes that arise from chemical and biological discovery campaigns. Clinical metabolomics studies provide one such testing ground: an interest in metabolites across a wide range of concentrations—where isolation and elucidation by NMR are usually unrealistic—makes LC-MS/MS followed by computational analysis a well-suited technique for the discovery of novel metabolites.

We first apply ICEBERG to the investigation of differentially enriched plasma metabolomic features from a post-traumatic stress disorder (PTSD) distress study collected from the Nurses’ Health Study cohort [54–56] (Fig. 3a). Earlier research has established a metabolite “distress” score based on features in the cohort associated with anxiety and depression phenotypes; this study identified other metabolite features that were associated with persistent PTSD symptoms. Previously conducted cohort statistical analysis identified a feature (198.1238 *m/z*) to be significantly negatively associated with persistent PTSD symptoms (*p* = 0.014) [56]. To elucidate this metabolite, we first infer the formula based on the MS1 and MS2 information, filtering a set of candidate formulae proposed by SIRIUS [59]. We then query isomers from PubChem with chemical formulae matching that of the most plausible formula, C_9_H_15_N_3_O_2_, from which we identify 8,413 candidates.

**Fig. 3:**
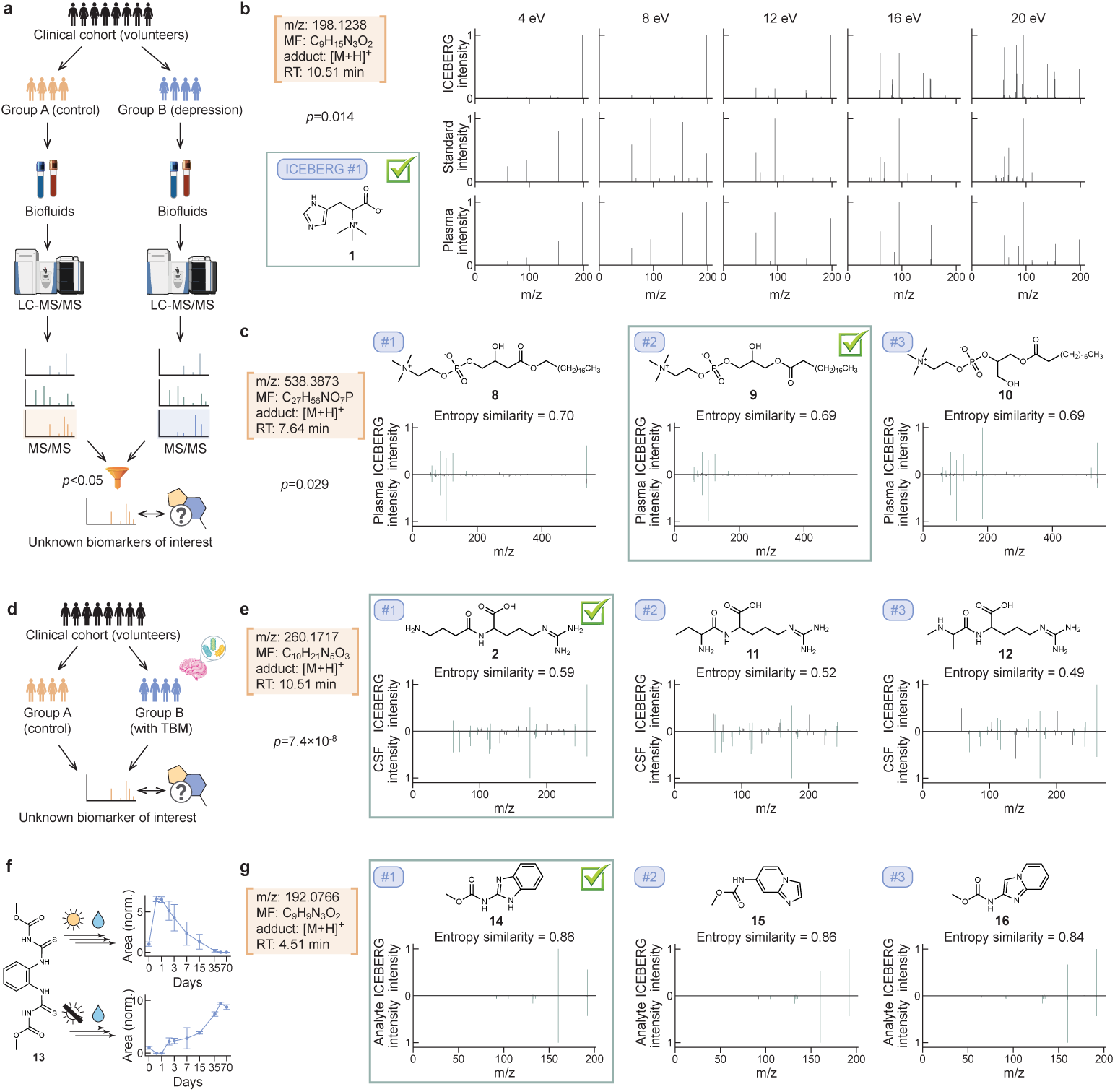
ICEBERG helps to elucidate novel structures in untargeted metabolomics and environmental studies with candidates from PubChem (**Fig. 1c**). Each structure annotation in this figure was verified by both retention time and MS/MS matching using an authentic standard. a, From the Nurses’ Health Study [54–56], a metabolomics study is performed for groups with or without depression, whereby statistically significant MS/MS features are identified. b, ICEBERG ranks hercynine (1) as the top-1 candidate for a biomarker that is negatively associated with depression. Alongside matches in RT, ICEBERG spectra predictions, MS/MS of the standard, and MS/MS from plasma align well across the five collision energies. Despite the difference in peak intensities, all three MS/MS exhibit the same trend of leftward peak-shifting across collision energies. c, Following ICEBERG predictions, LPC 19:0 (9) is identified as a novel biomarker positively associated with depression. ICEBERG predicts similar fragmentation patterns for the top-3 structures, where 9 is the most biologically plausible. LPC: lysophosphatidylcholine. d, A clinical study of tuberculous meningitis (TBM) [57, 58] yields metabolomic signatures of potential biomarkers of interest. e, ICEBERG’s top-ranked prediction, GABA-Arg (2), is verified as an unknown feature found to be negatively correlated with TBM infection and negatively correlated with mortality rate. GABA: gamma-aminobutyric acid. f, Thiophanate methyl (13) is a widely used fungicide. Under controlled abiotic degradation, an unknown degradant is found to accumulate in dark samples but is transient in light samples. g, ICEBERG ranks carbendazim (14) as the top-1 structure, which is further confirmed by an authentic standard.

ICEBERG is used to simulate the MS/MS spectra of these candidate structures with experimental parameters: an [M+H]^+^ adduct and collision energies of 4, 8, 12, 16, 20 eV, which correspond to 10%, 20%, 30%, 40%, 50% normalized collision energies (NCEs), respectively. Spectral entropy similarities are computed between each candidate *in silico* MS/MS and the true experimental spectrum, averaged among all collision energies. This retrieval strategy proposes hercynine (**1**) as the top-ranked structure (Fig. 3b). Following ICEBERG’s prediction, a hercynine standard was ordered, experimentally analyzed, and confirmed as a match by both RT and MS/MS (Extended Data Fig. 5a). Hercynine is a histidine derivative with known correlations with fungi or mushroom food intake [60, 61] but limited documented correlation with psychiatric state. The related derivative ergothioneine is well-established as an antioxidant and known to have neuroprotective effects [62], leaving room for further exploration and validation of possible causal biological mechanisms.

The same trauma study identified another higher-mass unknown feature (538.3873 *m/z*) that is positively associated with persistent PTSD symptoms (*p* = 0.029) [56]. 55 candidates with the formula proposed by SIRIUS, C_27_H_56_NO_7_P, are retrieved from PubChem. We use ICEBERG to simulate [M+H]^+^ spectra for these 55 candidates at the same five NCEs (Fig. 3c). The second-ranked structure **9** contains a canonical glycerol subgroup distinctive of lysophosphatidylcholine (LPC) and lysophosphatidylethanolamine (LPE) lipids, a subgroup not present in similarly-scored structures **8** and **10**. Based on this signature and class-level associations, we ordered LPC 19:0 (**9**), and confirmed its structure by RT and MS/MS matching to the standard (Extended Data Fig. 5b). Alongside several glycerophospholipids have been previously found to be implicated in mental disorders [63] and neurodegenerative disorders [64], LPC 19:0 is a novel biomarker of persistent PTSD symptoms; the abundance of metabolites in the same glycerophospholipid class, such as LPE 18:0 and LPC 34:2, was previously found to be statistically significant by Zhu et al. [56].

We would like to note that the results reported above involves human candidate selection based on ICEBERG predictions. There are another three candidates that were selected, purchased, but not matched in RT. More details can be found in Supplementary Section S5.

### Elucidation of biomarkers in clinical metabolomics studies of tuberculous meningitis

Following the same elucidation approach, we next apply ICEBERG to a tuberculous meningitis (TBM) study comprising 1,069 patients from Southeast Asia, from whom both cerebrospinal fluid (CSF) and plasma metabolomic samples were collected [57, 58, 65] (Fig. 3d). This study yielded an unknown feature (260.1717 *m/z*) negatively correlated with both TBM infection (*β* = −2.844*, p* = 7.4 × 10^−8^) and TBM mortality (*β* = −1.24*, p* = 0.07) [58]. We consider 61 candidates collected from PubChem with formula C_10_H_21_N_5_O_3_ and generate their [M+H]^+^ spectra at 10%, 20%, 30%, 40%, 50% NCEs. ICEBERG ranks GABA-Arg (**2**), an arginine conjugate of the GABA (gamma-aminobutyric acid) neurotransmitter [66], as the top structure (Fig. 3e). Several other arginine conjugates yield similar spectral similarity scores, but the gamma-butyric substructure is most structurally plausible as measured by predicted spectral similarity to the experimental CSF spectrum. A custom-synthesized authentic standard of GABA-Arg exhibited an exact RT and MS/MS match, validating ICEBERG’s proposed annotation (Extended Data Fig. 5c). ICEBERG has sufficient structural sensitivity to distinguish between subtly different isomers **2**, **11**, and **12** and correctly identify this feature as GABA-Arg.

While other GABA-amino acid conjugates have been identified and characterized, such as homocarnosine, a GABA-histidine conjugate thought to confer neuroprotective benefits following ischemic stroke [67], few other GABA conjugates with proteogenic amino acids have recorded biological significance to date.

### Elucidation of an abiotic degradant of thiophanate methyl pesticide

The importance of structural elucidation extends beyond metabolomics and into fields including environmental forensics. Here, using ICEBERG, we aim to characterize novel hydrolysis and photolysis products of pesticides collected in a controlled degradation study. Aqueous solutions of thiophanate methyl (**13**), a heavily used pesticide, were aged in simulated sunlight and dark conditions under environmentally relevant starting concentrations of 10 parts-per-billion (ppb). Aliquots were collected between 0 and 70 days to capture time-resolved system dynamics. Experimental features were collected using an untargeted, data-dependent acquisition mode with three stepped collision energies at 30%, 40%, and 60% NCEs. The challenge of identifying low parts-per-billion compounds and mapping their largely unexplored reactivity makes environmental forensics a compelling application for LC-MS/MS elucidation [7].

Thiophanate methyl (**13**) yielded an unknown (i.e., unmatched to reference spectra) degradation feature (192.0766 *m/z*) in both light-exposed and dark conditions but with substantially different temporal dynamics (Fig. 3f). To determine its identity, we employ the same formula candidate retrieval strategy, retrieving 4,790 C_9_N_9_N_3_O_2_ structures from PubChem and using ICEBERG to simulate all isomers’ spectra at the three collision energies. Visual inspection of the top-ranking candidates illustrates their close structural similarity, differing only in ring connectivity and subsequent spectral similarity; yet, the benzene ring from **13** is only preserved in the top-1 candidate, carbendazim (**14**) (Fig. 3g). Acquisition and analysis of an authentic standard confirmed a match based on both RT and MS/MS (Extended Data Fig. 5d).

Carbendazim is a known metabolite, a known photolysis product of thiophanate methyl (**13**) [68–71], an active fungicide, and a known pollutant in environmental samples [72]. Carbendazim is also well-classified as a reproductive toxin and carcinogen to both aquatic life and humans, with concentrations as low as 12.4-1240 ppb considered to confer an increased cancer risk [73–75]. However, carbendazim has not thus far been detected as a hydrolysis product of **13**. The kinetic patterns are distinct under light and dark conditions, where **14** is transient in light samples but accumulates in dark samples. The observed behavior may indicate the potential for carbendazim to accumulate in waters outside of the photic zone, in sediments, and within the human body, where photolytic pathways are inaccessible.

### Elucidation of isobaric reaction products facilitates data-rich experimentation

Beyond the detection of compounds in biological and environmental samples, we further demonstrate the utility of ICEBERG in synthetic chemistry. We report a proof-of-concept in assessing the relative reactivity of substrates through pooled experimentation. Traditional reaction screening evaluates reactants in a singleton fashion, in parallel or in sequence, which can be time-consuming and labor-intensive depending on the number of combinations under consideration (Fig. 4a). In contrast, pooled screening enables a single experiment to test multiple substrates (Fig. 4b). However, these approaches typically use MS1 to deconvolute (pooled) reaction products with unknown retention times, which imposes the critical limitation that products must be non-isobaric and therefore easily distinguishable [14, 77, 78]. Here, we show ICEBERG can enable pooled reaction screening in settings even with isobaric substrates (and thus expected isobaric products) (Fig. 4c), expanding the utility of these data-rich experimental workflows. This use case is distinct from the prior demonstrations in that the analytical challenge is not to identify fully unknown products, but rather to disambiguate expected products from a mixture where retention times are not known and no authentic standards are available.

**Fig. 4:**
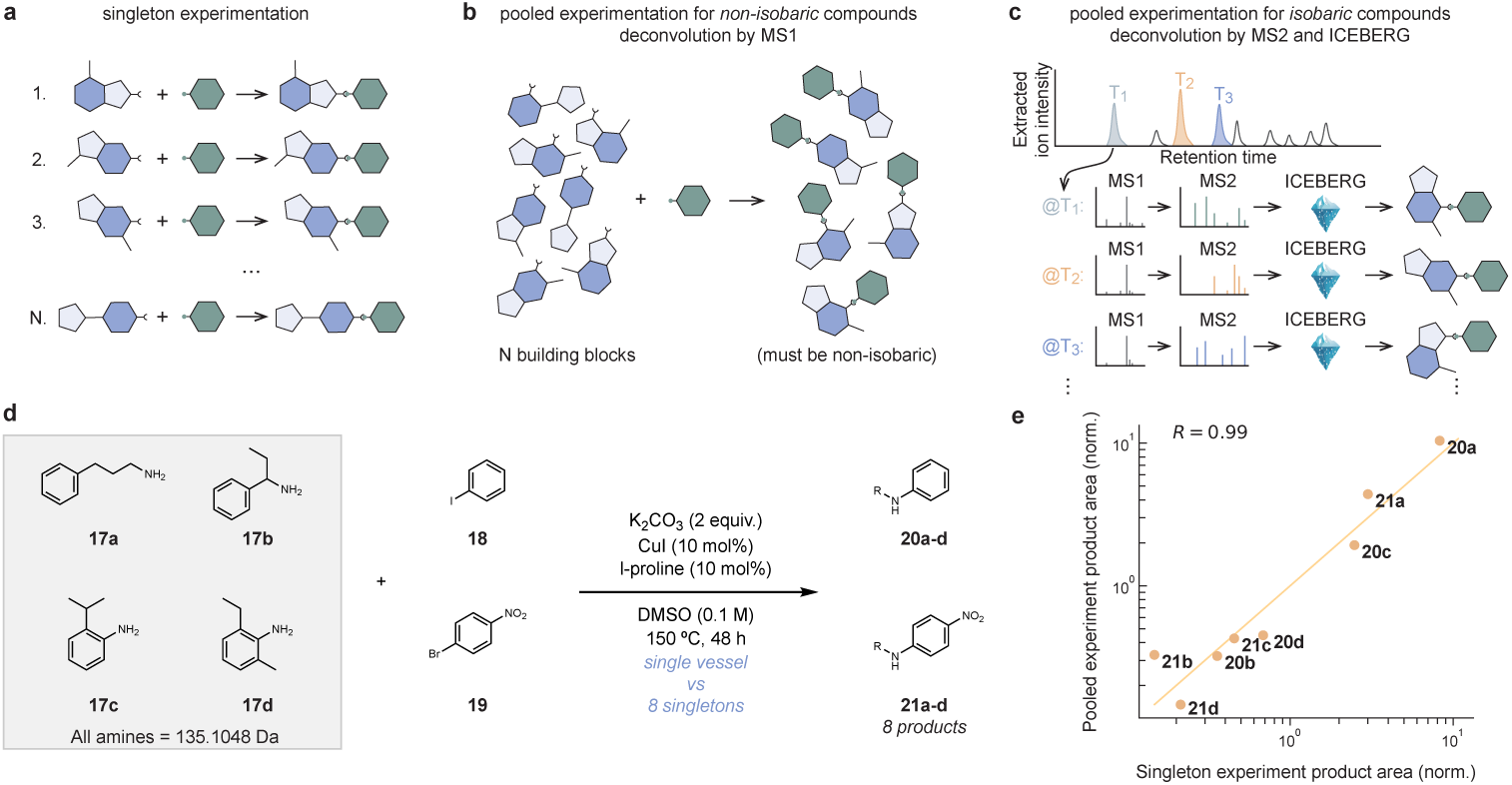
ICEBERG enables the deconvolution of pooled reaction screening experiments where candidates are derived from virtual enumeration (**Fig. 1d**). a, Traditional singleton experimentation requires running N experiments for N starting materials, which is labor-intensive. b, Pooled experimentation is more scalable, where a single pooled reaction is enough to test all N building blocks. However, a key limitation hindering the applicability is that pooled experimentation requires all potential products to be non-isobaric. c, In this work, LC-MS/MS and ICEBERG enable the deconvolution of isobaric products, allowing more flexibility in experimental design. d, As a validation case study, we use four isobaric anilines (17a-17d) and run C-N coupling [76] with iodobenzene (18) and 1-bromo-4-nitrobenzene (19). Such a pooled reaction creates two batches of isobaric products, which are further distinguished by LC-MS/MS and ICEBERG. e, The abundance of products in the pooled reaction aligns well with singleton experiments, as measured by the normalized extracted ion chromatogram area. The yellow line is the parity line (i.e., *y* = *x*).

We exemplify our MS/MS-enabled workflow with the analysis of reactivity trends across eight copper-catalyzed C–N coupling reactions. Using conditions reported by Zhang et al. [76], four isobaric amines were reacted with two halides within the same reaction vessel (Fig. 4d). Eight MS/MS features with the expected precursor masses (by MS1) were identified in quantifiable abundance after 48 hours at 150 °C. The eight products were matched to the eight features by the maximal linear assignment based on all pairwise spectral entropy similarities (Extended Data Fig. 6). It is worth noting that ICEBERG did not always rank the correct structure at the top 1, but a joint optimization of all spectrum-structure annotations yielded the correct assignment. In this study, the expected products **20c**and **20d** co-elute, but we are still able to distinguish them using a diagnostic fragment at 183.1043 *m/z* that ICEBERG recognizes would only appear in **20d**(Extended Data Fig. 7b). Each product is quantified by integrated extracted ion chromatogram (XIC) normalized using an internal standard (IS).

The intention of pooled screening is to provide an efficient way of accessing data-rich information. We therefore validate the pooled results by comparing normalized product abundances to the results of eight singleton experiments. All structure assignments are confirmed by RT (Extended Data Fig. 7a) and MS/MS matches. We also confirm that the reactivity trends among the four isobaric amines are largely preserved when comparing the pooled and singleton results (Fig. 4e). Though applied here to only four isobaric amines, these results illustrate how ICEBERG can enable deconvolution of isobaric and even co-eluting products that would otherwise impede such pooled reaction screening campaigns. We acknowledge the potential limitation of not being able to perfectly deconvolute isobaric products if the pool becomes larger or is comprised of structures that fragment similarly. To mitigate this limitation, one can predict whether two products will be easily deconvoluted by MS2 by fragmenting them *in silico* when designing the pool.

### Elucidation of biosynthetic pathways in *Withania somnifera*

Another elucidation strategy arises in applications such as analyzing biosynthetic pathways. To find close structural candidates, we can leverage the alignment strategies used in molecular networking for matching unknown MS/MS spectra to known reference spectra [79]. We further build upon these alignments to address the computational problem of localizing the specific site on the known molecule where the unknown structure differs as recently formalized by the ModiFinder algorithm [80]. Instead of utilizing combinatorial fragmentation explanations for MS/MS peaks, as with ModiFinder, we hypothesized that the use of ICEBERG-generated spectra and substructural explanations could localize where structural changes had taken place. This non-retrieval workflow, uniquely enabled by ICEBERG’s explainability, is a proof-of-concept that helps address the setting where hypothetical structures would be difficult or impossible to differentiate by MS/MS spectra alone, making retrieval-based identification infeasible.

We apply ICEBERG to investigate the biosynthetic pathway in *Withania somnifera* that produces bioactive compounds withanolide A (**30**) and withaferin A (**31**), among other withanolides, from 24-methylene cholesterol (**22**). Purified withanolides exhibit anti-cancer, anti-inflammatory, cardioprotective, neurological, and immunomodulatory activities [81–84] and clinical trials demonstrate that *W. somnifera*extracts can reduce depression, anxiety, and insomnia [85, 86]; improve cognitive function and memory [87]; and enhance cardiorespiratory endurance and recovery from stroke [88, 89]. Elucidating the enzymes involved in withanolide biosynthesis might enable biomanufacturing of withanolides for clinical and pharmaceutical applications. Until recently, only the first committed step of withanolide biosynthesis was known: the conversion of 24-methylene cholesterol (**22**) to 24-methyl desmosterol (**23**) by the sterol isomerase 24ISO [90]. To elucidate additional steps, candidate genes for involvement in withanolide biosynthesis were identified by Reynolds et al. [91] through biosynthetic gene cluster mining. Candidate genes were tested by heterologous expression in *Saccharomyces cerevisiae* (yeast) followed by metabolite extraction and LC-MS/MS analysis to identify pathway intermediates. Further experimental details are reported in ref. [91]. For each reaction, the mass difference and MS/MS spectra of the reactant and the product are known.

We use ICEBERG to predict potential modification sites for six intermediates using their respective known reactants as reference MS/MS spectra (Fig. 5a). For instance, in the transformation from compound **26** to **27**, we analyze the experimental spectrum of **27** against the experimental spectrum of **26** to identify peak pairs differing by 15.999 Da, indicative of hydroxylation. The ICEBERG-predicted peak annotations then provide the candidate substructures responsible for those peaks. We score each reference structure atom by summing intensities from matched peaks where its substructure includes that atom; atoms scored over 0.8 are proposed as probable modification sites. In the case of **26** → **27**, we identify six probable hydroxylation sites. In contrast, using the previously demonstrated retrieval pipeline can not distinguish the true structure from other isomers because certain substructures do not further fragment in MS/MS, making it challenging to locate the exact hydroxylation site. Also, other retrieval-based structural elucidation tools such as SIRIUS [37] or MIST [35] do not support this workflow as it would require an *in silico* fragmentation model.

**Fig. 5:**
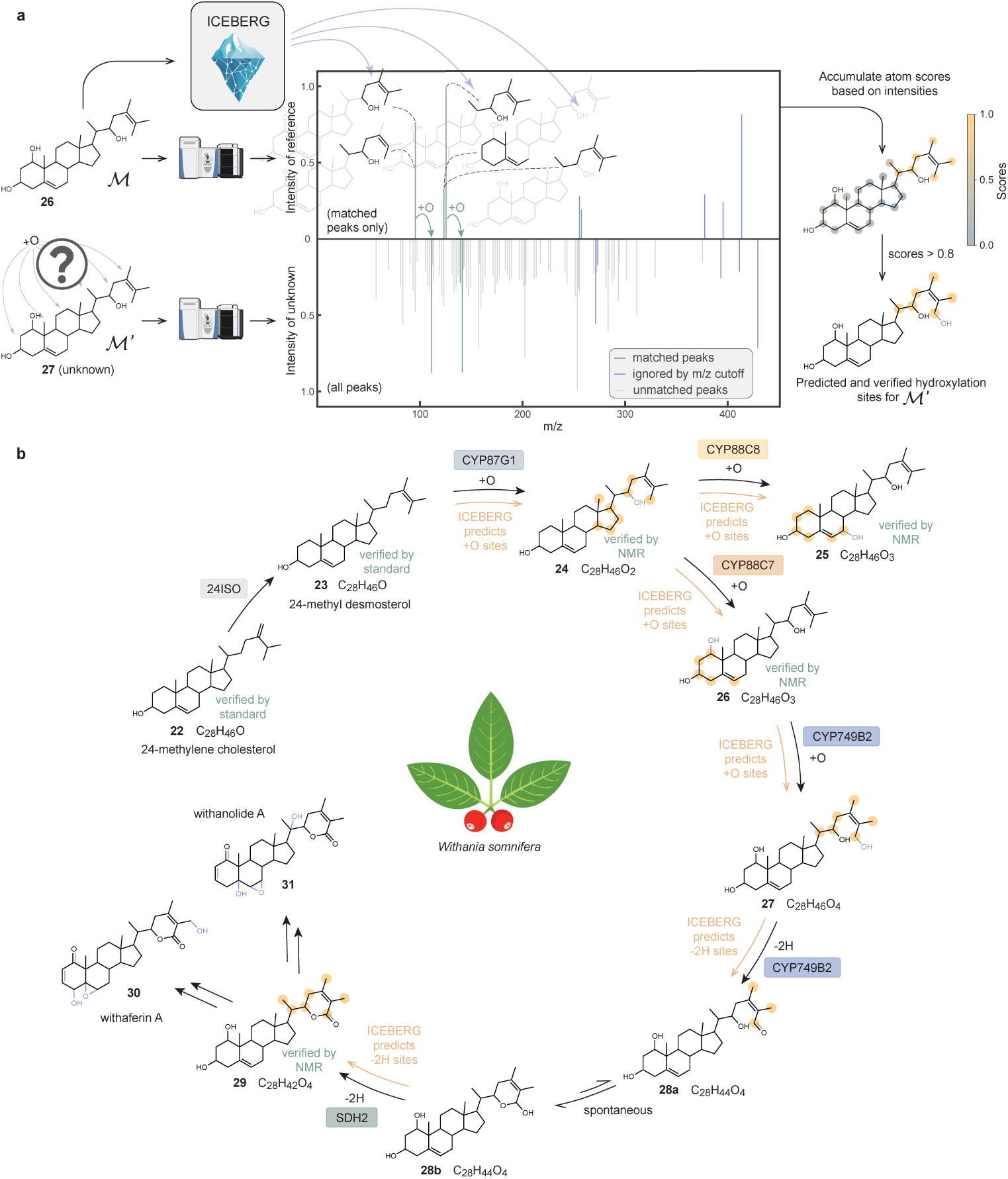
ICEBERG facilitates the elucidation of enzymatic function in a biosynthetic pathway with candidates derived from close reference structures (**Fig. 1d**). a, ICEBERG identifies high probability modification sites based on comparing MS/MS spectra for a known and an unknown structure. In this example, we aim to predict the hydroxylation site of 26 as a precursor to the unknown 27. Pairs of peaks with a mass shift of +O (15.9997 Da) are matched across the two spectra and annotated with the ICEBERG-predicted substructures that explain them in the reference structure (shown in black). Aggregation of intensities from each peak yields a set of predicted modification sites based on the resulting per-atom scores. Peaks over a cutoff of 200 *m/z* are ignored to focus on smaller fragments that can better localize information for structural elucidation. Sites with scores *>* 0.8 are shown in yellow, while the verified modification site is in green. b, This elucidation strategy is applied to a multi-step enzymatic pathway of *W. somnifera* that converts 24-methylene cholesterol (22) into bioactive compounds 30 and 31. ICEBERG successfully suggests potential modification sites for putative intermediate structures (24, 25, 26, 27, 28, 29) in the pathway by taking their precursor structures as references. Confirmed modifications are annotated in green, as verified by either NMR or inferred from other structures in the pathway with high confidence. With the biosynthetic steps to 30 and 31 yet to be resolved, we annotate all known net structural modifications relative to 29 in blue.

While ICEBERG does not propose a unique structure for each pathway intermediate in this setting, it narrows down potential sites of modification to streamline downstream metabolite identification. Viewed as a classification problem, this process achieves perfect recall: the true modification site is always contained within the 3-8 proposed candidates. This refinement is especially valuable given that NMR validation requires both bulk production and labor-intensive purification. With this stepwise elucidation strategy with ICEBERG, the biosynthetic pathway from compound **23** to **29** is successfully reconstructed (Fig. 5b), with key intermediates confirmed by NMR as shown in Supplementary Section S10. Altogether, these results demonstrate ICEBERG’s ability to guide and ultimately accelerate the structural elucidation of biosynthetic intermediates in complex natural product pathways.

## Discussion

In this work, we report a geometric deep learning framework, ICEBERG, that learns to simulate fragmentation in tandem mass spectrometry with state-of-the-art accuracy. We demonstrate successful applications of ICEBERG in identifying clinical biomarkers associated with depression (**1**, **9**) and tuberculous meningitis (**2**), elucidating an abiotic degradant of thiophanate methyl pesticide (**14**), enabling pooled reaction screening by deconvoluting isobaric reaction products (**20a-d**, **21a-d**), and annotating intermediate products in biosynthetic pathways in *W. somnifera* (**24-29**). These successful applications demonstrate structural elucidation in diverse chemical, biological, and medical domains and underscore the promise that ICEBERG-aided elucidation workflows with LC-MS/MS hold for the broader scientific community.

We envision several key avenues for future improvement that may enhance the utility of ICEBERG in additional prospective elucidation campaigns. First, ICEBERG accounts for rearrangements through its fragmentation prediction and allowance of ±6 hydrogen shifts; while this provides coverage of most known rearrangement rules (Supplementary Figures S1 and S2), revisions to the architecture might yield a more general solution. Second, we have focused our demonstrations on settings where structural hypotheses can be derived from PubChem, reaction enumeration, or minor modifications relative to a reference structure. There are many cases where structural candidates are not available, motivating the use of *de novo* generation approaches [38–42]. ICEBERG can be used to generate virtually unlimited numbers of simulated spectra to enable large-scale training of these generative models as foundation models for structure elucidation. Finally, as a practical consideration, ICEBERG could be augmented with a confidence estimator to help scientists prioritize which authentic standards to synthesize or purchase for unknown spectral features of interest.

We envision that deep learning-based MS/MS prediction will play a pivotal role in structural elucidation workflows across the community. The advances in ICEBERG presented here demonstrate a leap in prediction performance that enhances the viability of using MS/MS prediction for structure elucidation. By releasing a capable model that is open source and readily accessible by the community, ICEBERG is able to be used by the community and integrated into existing and newly developed structure elucidation pipelines, which in turn will drive new discoveries both scientifically and algorithmically.

## Methods

### Molecular space

Within the scope of this paper, molecules consisting of the elements C, H, N, O, S, P, F, Cl, Br, I, Si, B, Se, Fe, Co, As, Na, and K with molecular masses below 1,500 Dalton are considered. Molecules beyond this scope are not supported by ICEBERG’s model design.

### ICEBERG model architecture

ICEBERG is a neural MS/MS predictor for small molecules. Traditionally, MS/MS predictors rely on combinatorial enumeration of bond breakages to mimic physical fragmentation processes [25, 92]. While physically grounded, these methods are computationally intensive and often inaccurate due to overestimation of intensities. Deep learning-based predictors significantly reduce inference time while maintaining competitive accuracy, although they either fully discard physical constraints [29, 93] or only partially retain them via subformulae [27, 28]. ICEBERG strikes a balance by generating a pruned fragmentation graph and predicting intensity values using transformers [94]. It is a two-stage model consisting of ICEBERG-Generate and ICEBERG-Score (Fig. 1a and Extended Data Fig. 1).

### ICEBERG-Generate

(Extended Data Fig. 1a). ICEBERG-Generate simulates high-energy collision-induced dissociation (HCD) by iteratively predicting the most probable fragments through stepwise atom and bond removals from the molecular graph. This autoregressive process supports multiple fragmentation steps, producing a directed acyclic graph (DAG) with the original molecule as the root and its fragments as children. At each step, the model receives a (sub)fragment as input and outputs the probabilities of atom breakages, which are then used as inputs for the next step. As the ground-truth data provides only *m/z* values and intensities—without annotated fragmentation graphs—we use the MAGMa algorithm [24] to generate DAG labels (please refer to Training ICEBERG for details). We allow up to three atom-breaking steps and a maximum of six bond breakages. All predicted fragments are ranked by their cumulative probabilities, and the top 100 are passed to ICEBERG-Score.

### ICEBERG-Score

(Extended Data Fig. 1b). ICEBERG-Score processes the fragmentation DAG to predict peak intensities for each fragment. The architecture comprises: 1) a GNN module that ingests molecular graphs and instrumental parameters to generate embeddings for both fragments and the root molecule, and 2) a set transformer [94] that predicts the fragment intensities. Multiple intensity values are predicted for the same fragment to account for hydrogen shifts and charge migration. Predicted intensities are merged into a mass spectrum using the corresponding *m/z* values. For more details, readers are referred to Goldman et al. [46] and the open-source code at https://github.com/coleygroup/ms-pred.

### Incorporating collision energy

Collision energy is a critical LC-MS/MS parameter that heavily influences fragmentation patterns. In Goldman et al. [46], spectra with varying collision energies were merged and treated as a single prediction task.

Here, we explicitly incorporate collision energy (in eV) using a 64-dimensional positional encoding (32 sine, 32 cosine terms, base 10,000) to encode its continuous value. We choose position encoding because it turns an arbitrary large value into a fixed-length vector, without sacrificing the granularity of differentiating as small as 1 eV. This encoding is concatenated with the input features for both ICEBERG-Generate and ICEBERG-Score.

### Incorporating negative mode and featurization

Earlier versions of ICE-BERG supported only positive mode adducts, such as [M+H]^+^, [M+Na]^+^, and [M−H_2_O+H]^+^. However, many metabolites (e.g., fatty acids, phosphorylated or halogenated compounds) are better fragmented in negative mode [95, 96], particularly [M−H]^−^. We incorporate negative mode by encoding new adduct types as one-hot vectors. Additional chemical knowledge is encoded as multi-hot vectors, including features such as positive/negative mode, alkali metal adducts, proton gain/loss, halogen adducts, loss of H_2_O or CO_2_, and gain of NH_3_. All these features are concatenated with the model input.

### Intensity prediction with charge migration

ICEBERG-Score predicts up to 13 × 2 = 26 intensity values per fragment. Part of ICEBERG-Generate’s output is the number of bond-breaking steps needed for each fragment, and we allow for a ±1 hydrogen shift for each step to accommodate rearrangements (Supplementary Figures S1 and S2). With a maximum of six bond breakages, this results in 13 hydrogen shift states (Extended Data Fig. 1b). We also incorporate both charge retention fragmentation (CRF) and charge migration fragmentation (CMF) [53]. For instance, for [M+A]^+^ and denoting B as the neutral loss, both [M−B+A]^+^ (CRF) and [M−B]^+^ (CMF) might be observed, i.e., CRF indicates adduct retention, whereas CMF indicates adduct detachment and charge migration. We introduce CMF as part of the technical contributions of this paper. While Goldman et al. [46] implicitly handled CMF for [M+H]^+^ with hydrogen shifts, other adducts such as [M+Na]^+^ or [M−H_2_O+H]^+^ require explicit modeling. The benefits of charge migration modeling are demonstrated in the ablation study (Extended Data Table 1).

### Dataset processing

The NIST’20 dataset [51] comprises standardized spectra collected using various instruments. Among these, HCD LC-MS/MS spectra—most commonly obtained via Orbitrap instruments—are predominant in structural elucidation. We therefore focus exclusively on HCD spectra. We assume that all deposited spectra have undergone basic noise filtering and perform no further preprocessing. We square root all intensities (between 0 and 1) to emphasize lower-intensity peaks. To align with our target use case, we exclude non-HCD spectra. Although this work does not include QTOF spectra, our approach can be extended to handle them by retraining or finetuning.

We aim to support the most common adduct types for the maximum applicability of ICEBERG. Due to the challenges in modeling, we exclude adducts with multiple charges (e.g., [M+2H]^2+^), multiple neutral molecules (e.g., [2M+H]^+^). Less abundant adducts that comprise less than 0.8% of the dataset (except [M+K]^+^) are also excluded. The following adducts are used for training and evaluation: [M+H]^+^, [M+Na]^+^, [M+K]^+^, [M−H_2_O+H]^+^, [M−2H_2_O+H]^+^, [M+NH_3_+H]^+^, [M−H]^−^, [M+Cl]^−^, [M−H_2_O−H]^−^, and [M−CO_2_−H]^−^. Collision energy values recorded as normalized collision energy (NCE) are converted to eV using:

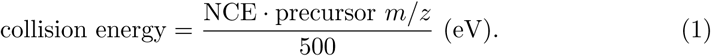

Molecules with unsupported atom types (Molecular space) and molecular masses larger than 1,500 Dalton are also filtered out. After filtering, the dataset contains 530,640 spectra across 25,541 unique molecules (Extended Data Table 2). All models use the same 8:1:1 train/validation/test split. We follow the data splits created by previous benchmarks [28, 46], while since these benchmarks are positive-adduct-only, we maintain the same split on positive adducts and build upon for structures that only appear in the negative mode.

### Training ICEBERG

#### Hardware setup

All experiments reported in this paper are conducted on a workstation with AMD 3995WX CPU, 4×NVIDIA A5000 GPU, and 512GB RAM.

#### Generating fragmentation DAG labels and training ICEBERG-Generate

ICEBERG-Generate requires annotated fragmentation paths, which are absent in raw NIST’20 data. We use the MAGMa algorithm [92] to enumerate all potential fragmentations. Each generated fragment is matched to a peak *m/z* value, allowing for ±6 hydrogen shifts and both [M−B+A]^+^ and [M−B]^+^ forms if the adduct is [M+A]^+^ (analogous rules apply in negative mode). High-resolution HCD spectra (i.e., Orbitrap, with 5 ppm accuracy) facilitate accurate fragment identification. For matched fragments, we construct a minimal spanning tree to serve as the fragmentation DAG label for training. ICEBERG-Generate is trained at a starting learning rate of 0.000996, decays by 0.7214 after every 5,000 steps, and has a dropout rate of 0.2. These hyperparameters are directly adapted from ref. [35]. Training for 16 epochs takes 6.2 hours on a single NVIDIA RTX A5000 GPU. Before training ICEBERG-Score, we need to use the pretrained ICEBERG-Generate to predict DAG labels on the training set, which takes 3.0 GPU hours.

#### Training ICEBERG-Score with spectral entropy loss

ICEBERG-Score is trained to minimize differences between predicted and reference spectra. Spectral entropy, as proposed by Li et al. [49], has been shown to outperform cosine similarity for spectrum comparison. While Goldman et al. [46] employed cosine loss, we adopt entropy loss to better emphasize low-intensity peaks:

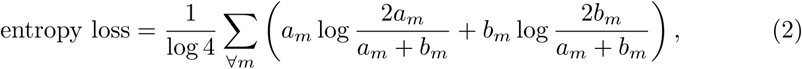

where *a_m_* and *b_m_* denote the normalized intensities at *m/z* = *m* in spectra 1 and 2, respectively. The normalization term log 4 reflects the maximum entropy divergence and ensures the entropy loss is between 0 and 1. Because many predicted intensity values are zero (due to absent fragments), intensity prediction resembles a classification task. Emphasizing low-intensity peaks aids the model in distinguishing relevant fragments. From a deep learning perspective, cross-entropy is also the natural choice for training classifier models, aligning with our adoption of the entropy-based loss. ICEBERG-Score is trained at a starting learning rate of 0.000736, decays by 0.825 after every 5,000 steps, and has a dropout rate of 0.1. Training for 21 epochs takes 9.2 hours on a single NVIDIA RTX A5000 GPU.

#### Contrastive finetuning

Contrastive finetuning has emerged as an effective regularization strategy to improve model generalization [47]. To enhance retrieval accuracy, we develop a contrastive finetuning procedure using negative samples from Pub-Chem and mutated molecules. Specifically, 50% of negative structures share the same chemical formula (from PubChem), and 50% are generated by single-step graph mutations [97, 98]. During training, ICEBERG predicts on one positive and three negative structures simultaneously, yielding the following pairwise distance matrix:

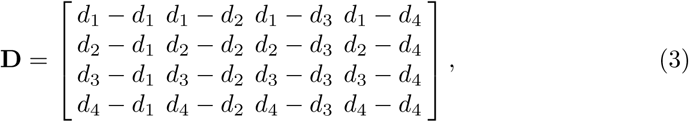

where *d_i_* is the entropy distance (Equation (2)) between the predicted spectrum for structure *i* and the ground-truth spectrum, and *d*_1_ is the positive sample. We encourage *d*_1_ to be the smallest via a differentiable soft ranking using the Sinkhorn algorithm [99, 100], where **D**^-^ = Sinkhorn(−**D***, τ*) and *d*^-^_1_ ∈ [0, 1] is the top-left entry of **D**^-^. Sinkhorn could be viewed as a soft, differentiable version of linear assignment, where a smaller temperature *τ* will lead to a more discrete output. We set *τ* = 0.05. When *d*_1_ is the smallest among {*d*_1_*, d*_2_*, d*_3_*, d*_4_}, Sinkhorn algorithm will yield a larger *d*^-^_1_. The larger *d*^-^_1_ is, the larger the gap between *d*_1_ and the next smallest *d_i_*(*i* ≠ 1). The contrastive loss is then:

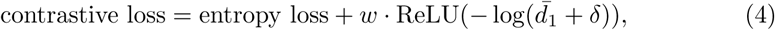

where *w* (empirically set to 1) scales the contrastive term, and the offset *δ* = 0.5 prevents overfitting by discouraging further maximization when *d*^-^_1_ is already large. To finetune ICEBERG-Score, we have to run ICEBERG-Generate on all decoy structures (10 per training sample), which takes 28.1 GPU hours. The finetuning step has a learning rate of 0.0005, decays by 0.825 after every 5,000 steps, and has a dropout rate of 0.1. Training for 10 epochs takes 7.9 hours on two NVIDIA RTX A5000 GPUs.

### Molecular formula identification

In these experimental applications, MS1 information is sufficient to infer the chemical formula with existing cheminformatic tools [59, 101, 102]. Candidate structures with consistent formulas, in this initial demonstration originating from PubChem, can then be narrowed down with ICEBERG. Matches are ultimately validated experimentally based on the chromatographic retention time and MS/MS spectrum of a pure reference compound (i.e., authentic standard).

We identify the molecular formula of unknown MS/MS features by either manual annotation or running SIRIUS [59]. For the structural elucidation campaign for GABA-Arg (**2**), we enumerate all possible chemical formula that matches its MS1 mass and manually check if it is a reasonable chemical formula by comparing MS2 peaks to the subformulae. For campaigns for hercynine (**1**), LPC 19:0 (**9**), and carbendazim (**14**), we run SIRIUS (6.0.6) under the command line interface and take the top formula prediction. For the reaction screening study (Fig. 4) and the elucidation of biosynthetic pathway for *W. somnifera* (Fig. 5), the formulae are derived based on known starting materials and reaction rules derived from the difference in *m/z*.

### PubChem candidates generation

We use our locally cached version of PubChem (downloaded in January 2024, with 116,124,942 SMILES) for all numerical evaluations and case studies. We compute chemical formulae for all structures, create a mapping from chemical formula to SMILES, and deduplicate SMILES and stereoisomers based on InChI keys. We also ignore salts, disjoint structures, molecules containing unsupported elements, and molecules that are heavier than 1,500 Dalton.

### Spectral similarity metric

We use spectral entropy similarity [49] as the metric to rank candidate structures for all structural elucidation campaigns reported in this work. For the elucidation of hercynine (**1**), LPC 19:0 (**9**), GABA-Arg (**2**), and carbendazim (**14**), their precursor ion peaks are masked, and the remaining peaks are normalized when computing the entropy similarity to avoid the bias introduced by the precursor peak. For the deconvolution of **20a-d** and **21a-d**, the precursor peak is retained because the relative peak intensity of the precursor dictates the disambiguation process. The experimental spectra are processed with electronic denoising [103], square-rooted to emphasize the lower-intensity peaks, and we only consider the top-20 peaks from each spectrum (except for withanolides that have rich fragmentation).

### Handling stepped collision energy

In the studies of the abiotic degradation of thiophanate methyl and pooled C-N coupling reaction screening, we ran LC-MS/MS at stepped collision energy to achieve rich fragmentation patterns with multiple energies while avoiding the time cost of running the same sample multiple times. In the stepped collision energy mode, the precursor ions are first fragmented at different energies, and then detected at the same time. To mimic the same process with ICEBERG, as the applied energy values are known (30%, 40%, 60% NCEs), we use ICEBERG to predict spectra at these energies, and then merge them to create the *in silico* spectra. When merging, spectra intensities are summed if their *m/z* values are the same to the 4th decimal point.

### Disambiguation of co-eluting peak areas

In the pooled screening of the C-N coupling reactions, **20c** and **20d** co-eluted, making it non-straightforward to measure the abundances of **20c** and **20d** in the pool because their peak areas are merged at the precursor mass. As predicted by ICEBERG, we use their diagnostic peaks in MS2, which are also in-source fragments in MS1, to compute their relative abundance. More specifically, we look for the XIC peak of 170.0965 *m/z* at the same RT as **20c**’s precursor peak, and 183.1043 *m/z* for **20d** (Extended Data Fig. 7b). We compute these two XIC peak areas (**20c**’s 170.0965 *m/z* PROD/IS: 8.19×10^−3^; **20d**’s 183.1043 *m/z* PROD/IS: 1.91×10^−3^) and use them as weights to split the merged precursor peak area. The merged XIC peak has PROD/IS: 2.378, and **20c**’s PROD/IS is estimated as 1.928, **20d**’s PROD/IS is estimated as 0.450.

### Structure annotation from reference structures

To annotate possible modification sites from a known reference, we extend the ModiFinder algorithmic framework [80] with ICEBERG. ModiFinder identifies possible modification sites from a known reference structure and a structurally similar unknown, where both of their MS/MS spectra are characterized. The original ModiFinder then utilizes a combinatorial fragmentation enumeration [92] to explain peaks with the *m/z* shift of interest, where intensity scores are accumulated for each atom. In this work, we replace the combinatorial enumeration with ICEBERG and introduce several technical adjustments. First, ICEBERG learns and predicts fragment structures, which are used to match peaks of interest by *m/z* values. If ICEBERG predicts multiple fragments with the same *m/z* value, all substructures are considered, and the score is reweighted by ICEBERG-predicted intensities for each of them. Second, we apply an *m/z* value cut-off (set to 200 for case studies in Fig. 5) to avoid accumulating scores for larger fragments, which contain less information to help locate the modification site. Finally, the scores are normalized to the range between 0 and 1, and candidates with atom-level scores *>* 0.8 are considered as possible modification sites, as highlighted in yellow in Fig. 5.

## Declarations

### Supplementary information

Supplementary Information is available for this paper.

## Acknowledgements.

The authors acknowledge Dr. Itai Levin, Mohammad Reza Shahneh, Jenna Fromer, Dr. Raji Balasubramanian, Dr. Sue Hankinson, Dr. Arjan van Larhooven, Dr. Reinout Van Crevel for assistance, and Dr. Guangqi Wu, Dr. Wenhao Gao, Dr. Keir Adams, Dr. Tianyi Jin, and Rui-Xi Wang for discussions.

## Ethics approval and consent to participate

Not applicable.

## Consent for publication

Not applicable.

## Data availability

The NIST’20 dataset used to train ICEBERG is a commercial resource and is available for purchase through several vendors worldwide https://chemdata.nist.gov/dokuwiki/doku.php?id=chemdata:distributors. Given the scale of effort required to purchase samples, run experiments, and collect the 530K spectra, and that NIST’20 is the only database where all spectra have collision energy annotations, this dataset is a reasonable investment in mass spectrum-related research in the absence of a thorough open-source replacement. We also provide the code to process raw data at https://github.com/rogerwwww/ms-data-parser. The LC-MS/MS data from case studies, DFT calculations, and spectra and SIRIUS project files for the MSnLib evaluation are deposited in MassIVE under the identifier MSV000097986.

## Materials availability

The vendor information of purchased analytical standards is presented in supplementary information (Supplementary Section S4).

## Code availability

All code for training and testing ICEBERG, together with example applications, is publicly available at https://github.com/coleygroup/ms-pred. As the NIST’20 dataset is a commercial resource requiring the purchase of a license, the pretrained weights will be made available to those with a license.

## Author contribution

R.W., M.M., S.G., and C.W.C. developed the ICEBERG model. R.W. and M.M. developed the protocol for real-world applications. B.M. and C.W.C. led the pooled C-N coupling reactivity study. J.A.-P., E.M., and C.B.C. led the LC-MS/MS analysis of clinical biomarkers. J.P. and D.L.P. led the pesticide degradation study. E.R. and J.-K.W. led the study of *W. Somnifera*. M.L. conducted the computational analysis of inductive collision energies. M.W. proposed the modification localization algorithm. All authors contributed to the writing of this paper.

---

**Extended Data Fig. 1:**
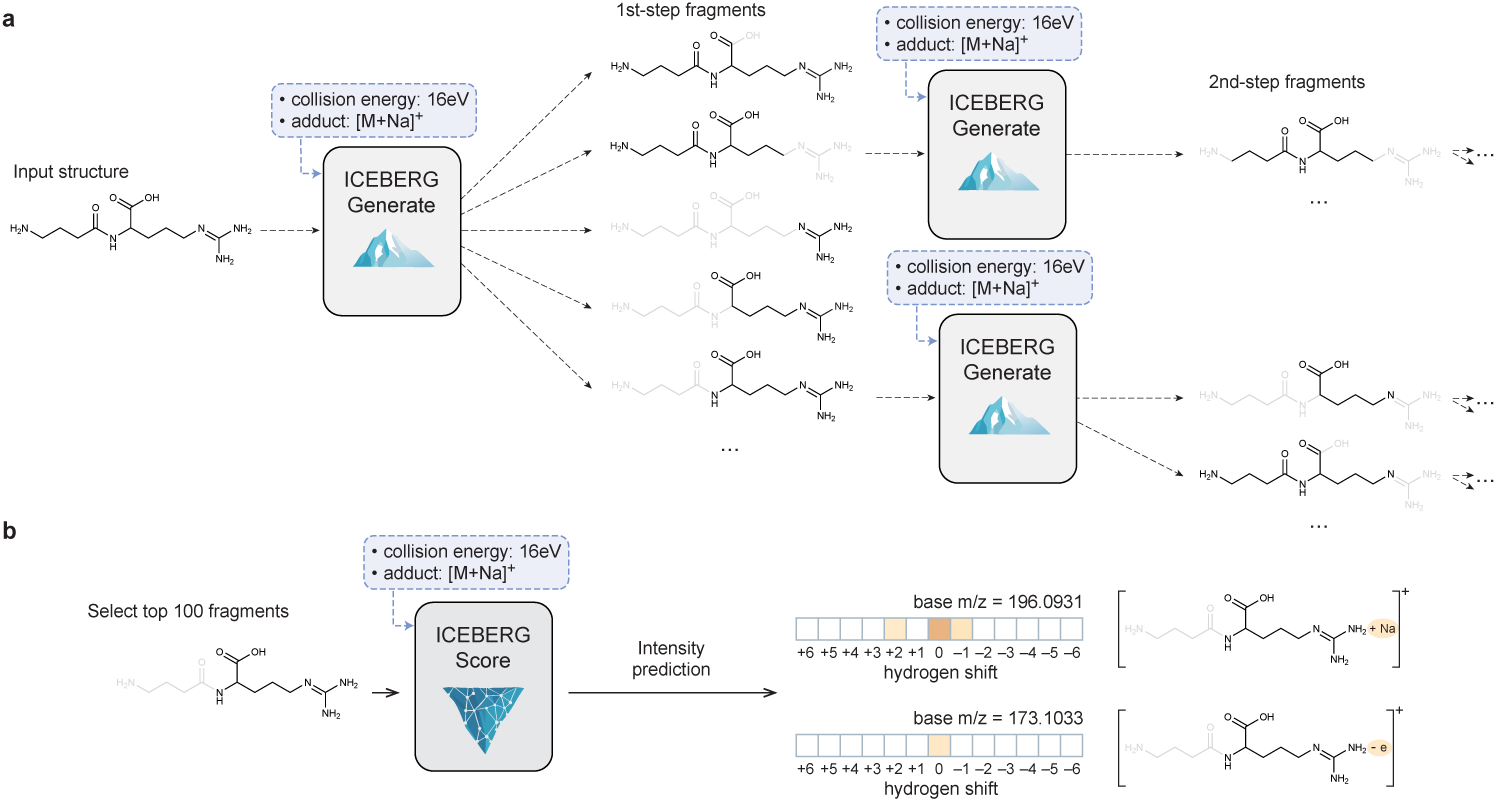
Details of ICEBERG-Generate and ICEBERG-Score with architectural improvements in this work. **a**, ICEBERG-Generate takes a (sub)structure as input and predicts plausible fragmentation. It is an auto-regressive model, i.e., the first-step fragments are fed into the model again to fragment them further, creating second-step fragments. The model predicts probabilities at each atom about whether this atom should be removed, and we allow up to 3 fragmentation steps for the fragmentation graph. **b**, ICEBERG-Score takes the top 100 fragments from the fragmentation graph and predicts intensities for each of them. The output dimension is 26, including up to ±6 hydrogen shift that accounts for rearrangements relative to two “base” *m/z* values that account for retaining or migrating the charge from the adduct ion (Na^+^ in this case).

**Extended Data Fig. 2:**
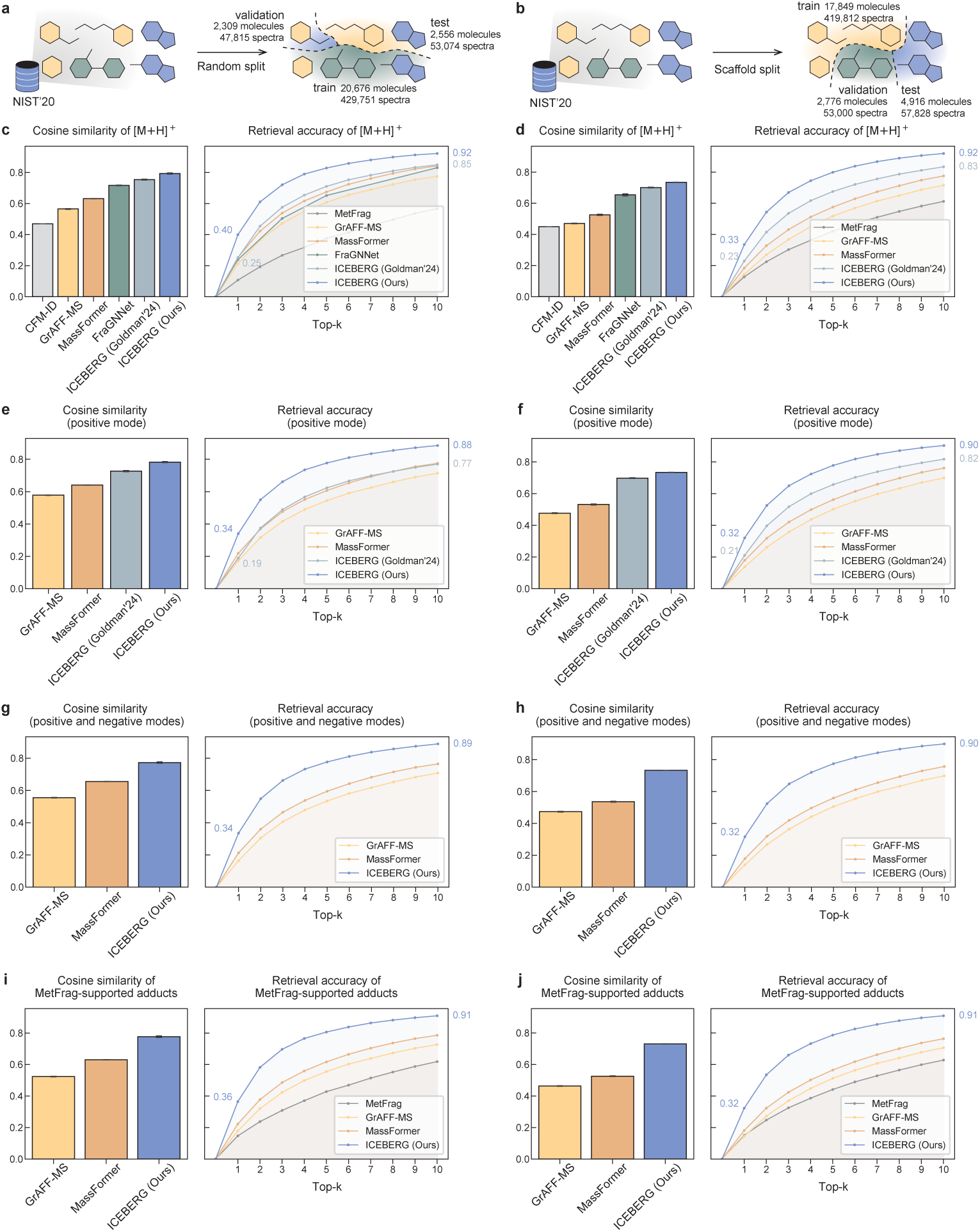
Quantitative evaluation results on different adduct types. Error bars with 95% confidence interval are reported for 3 random seeds for cosine similarity. Only mean retrieval accuracies are plotted, and their confidence intervals are reported in Supplementary Tables S2, S4, S6 and S7. **a**, The same random split as reported in Fig. 2 is used, and models are the same without retraining. **b**, We report results on a scaffold split, a more challenging setting where the models are tested on unseen molecular scaffolds. **c**, ICEBERG predicts spectra more accurately for the most common [M+H]^+^ adduct compared to other prediction tools. **d**, Similar trend is observed for [M+H]^+^ on the scaffold split, where ICEBERG outperforms peer methods by a significant margin. **e-f**, For all positive mode adducts ([M+H]^+^, [M+Na]^+^, [M+K]^+^, [M−H_2_O+H]^+^, [M−2H_2_O+H]^+^, [M+NH_3_+H]^+^) under both random and scaffold splits, ICEBERG outperforms all competing methods in terms of spectrum prediction accuracy and retrospective evaluation of retrieval accuracy. **g**, For all positive and negative adducts (aforementioned positive adducts and [M−H]^−^, [M+Cl]^−^, [M−H_2_O−H]^−^, [M−CO_2_−H]^−^), with random split, ICEBERG reaches state-of-the-art accuracy. Peer methods that do not support negative mode are excluded for comparison. **h**, When switched to scaffold split, ICEBERG still performs as the state-of-the-art. In terms of retrieval accuracy, the performance degeneration is not significant from random split to scaffold split, and from positive mode to negative mode. **i-j**, ICEBERG outperforms the learning-free fragmentation heuristic MetFrag among the adduct types supported by MetFrag ([M+H]^+^, [M+Na]^+^, [M+K]^+^, [M+NH_3_+H]^+^, [M−H]^−^, [M+Cl]^−^).

**Extended Data Fig. 3:**
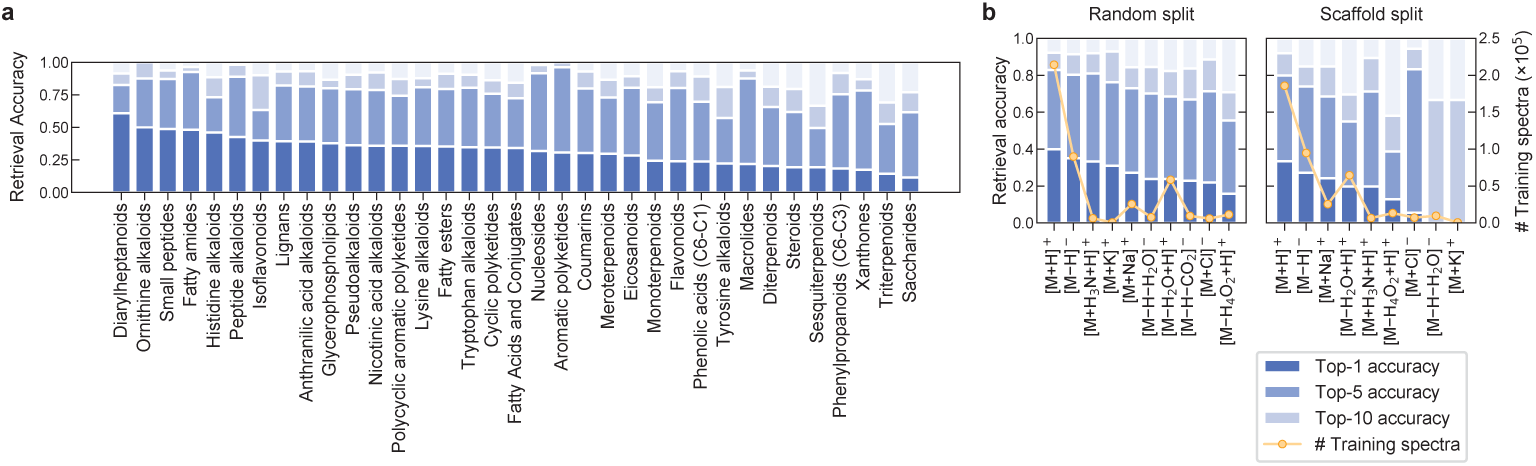
Extended quantitative evaluation results on the 20 benchmark. **a**, Retrieval accuracies differences on NIST’20 by compound class as assigned by NPClassifier [104]. Classes with >20 compounds are included for visualization. **b**, Retrieval accuracy on NIST’20 as a function of adduct type shows that [M+H]^+^ and [M−H]^−^ are the best-performing adduct types, consistent with these adducts comprising the plurality of data in NIST’20.

**Extended Data Fig. 4:**
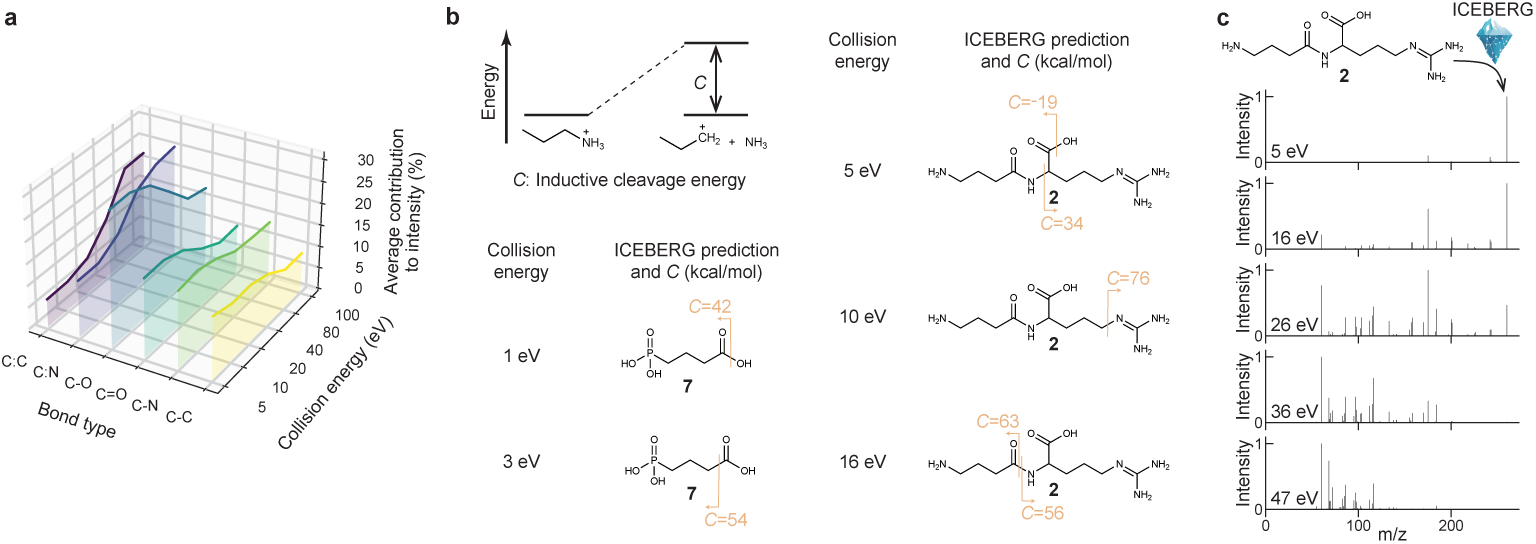
Qualitative evaluation demonstrates ICEBERG’s interpretability as matched by chemistry intuition and DFT calculations. **a**, For the six most abundant bond types in NIST’20 testing dataset, ICEBERG’s prediction of the average contribution to spectral intensity matches chemistry intuitions. Bond type notations follow SMARTS primitives, where C:C and C:N denote aromatic bonds, C=O denotes double bond, and C-O, C-N, C-C denote single bonds. C-O and C=O bond fragmentations are of greater relative importance at lower collision energies, and their trends are similar, alongside the double bond exhibits stronger. ICEBERG also learns that aromatic bonds are stable at lower collision energies, but break more often when the collision energy goes as high as 100 eV. **b**, ICEBERG learns bond cleavage orders that associate with DFT calculation of inductive cleavage energy (*C*). Inductive cleavage energy is defined as the bond dissociation energy for charged ions. In the case studies of **2** and **7**, ICEBERG predictions align with *C* in most cases, except for the outlier at 10 eV for **2**. **c**, A sweep of collision energies for **2** illustrates that ICEBERG learns the expected shift from higher *m/z* peaks to lower *m/z* peaks as collision energy increases.

**Extended Data Fig. 5:**
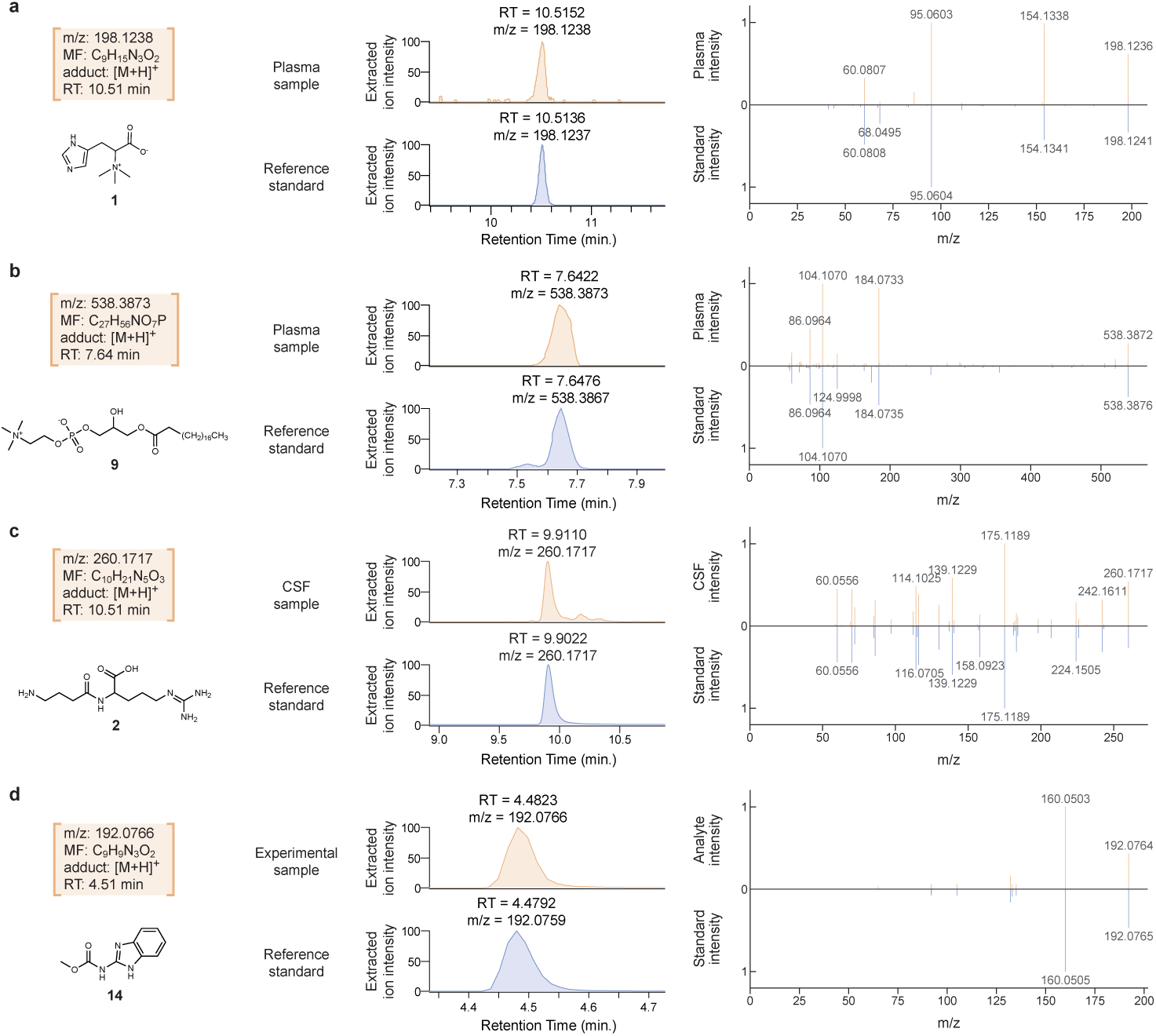
Matching of authentic standards and the biomarkers and pesticide degradant shows agreement in both RT and MS/MS fragmentation. For each elucidated structure, we purchase the reference standard, run it with the same LC-MS/MS instrument again, and verify the structure by matching both RT and MS/MS. Experimental spectra of the unknown and the standard are shown in the upper and lower halves of the mirror plots, respectively. Mirror plots viewable on MassIVE for each of these samples are available for hercycine (**1**), LPC:19-0 (**9**), GABA-Arg (**2**), and carbendazim (**14**), with embedded links in the PDF version.

**Extended Data Fig. 6:**
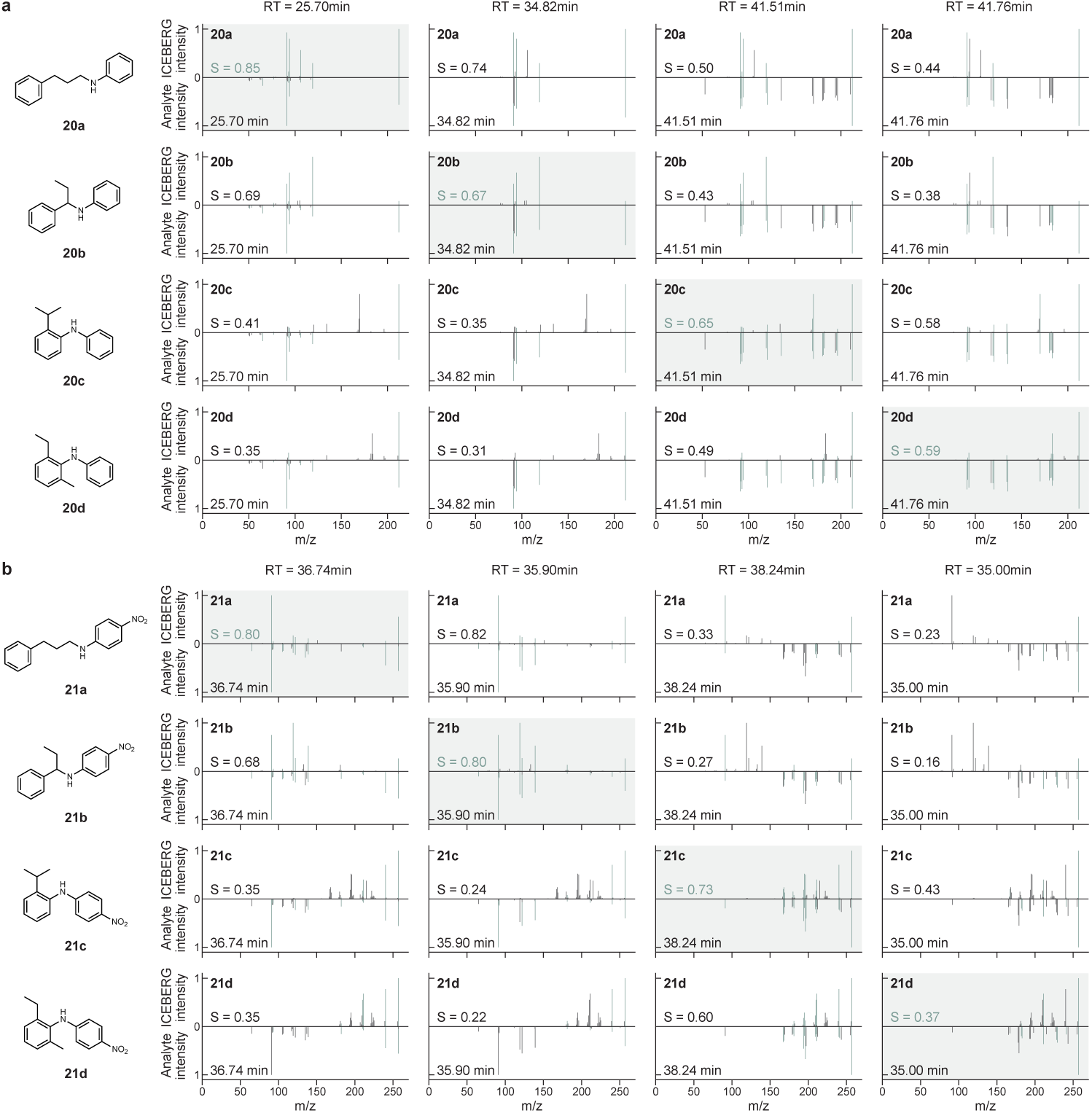
ICEBERG deconvolutes isobaric reaction products in pooled screening. ICEBERG successfully matches isobaric products to their MS/MS. In each mirror plot, the upper half is ICEBERG-predicted spectra (with structures at each row), and the lower half is experimental spectra (with RTs at the top of each column). S denotes the spectral entropy similarity. The assignment of structures to RTs is obtained by solving a maximal linear assignment problem from ICEBERG-predicted similarities. **a**, isobaric products and matching at 212.1432 *m/z* (**20a**-**20d**). **b**, isobaric products and matching at 257.1285 *m/z* (**21a**-**21d**).

**Extended Data Fig. 7:**
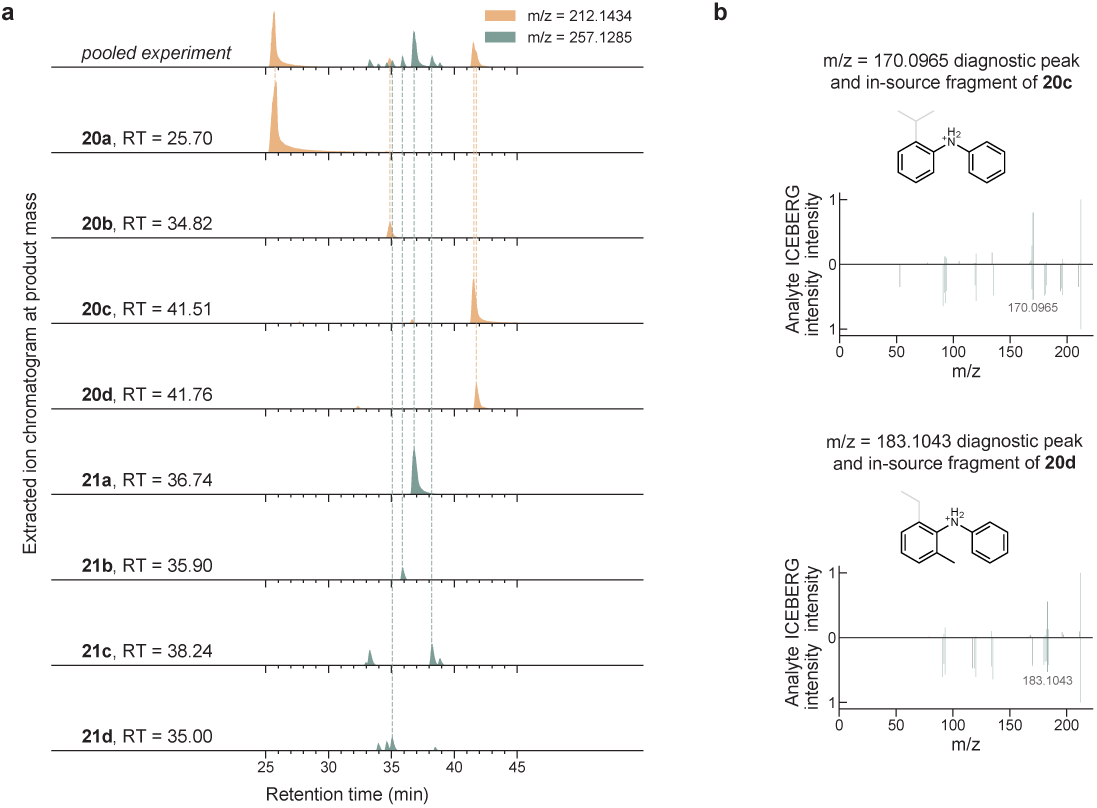
Comparison of singleton and pooled reactions validates structural elucidation of our C-N coupling products. **a**, Singleton experiments validate our structural elucidation of the products by RT and MS/MS match. The XIC peak intensities are normalized and square-rooted for better visibility of the lower intensity peaks. Mirror plots comparing pooled and singleton products are available via MassIVE for **20a**, **20b**, **20c**, **20d**, **21a**, **21b**, **21c**, **21d**, with embedded links in the PDF version. **b**, For **20c** and **20d** that are coeluting in the pooled experiment, they are deconvoluted by ICEBERG with the diagnostic peak at 183.1043 *m/z*. We visualize ICEBERG predictions with the first two MS2 scans from the pooled peak, representing **20c**, and the last two MS2 scans for **20d**. Since 183.1043 *m/z* and 170.0965 *m/z* are unique in-source fragments for **20c** and **20d**, respectively, the XIC peak areas of these in-source fragments are used to help calculate peak areas of **20c** and **20d** from the pooled experiment.

**Extended Data Table 1:** Ablation study of ICEBERG technical improvements.

| Top-$k$ accuracy | Top-1 | Top-2 | Top-3 | Top-4 | Top-5 | Top-6 | Top-7 | Top-8 | Top-9 | Top-10 |
| --- | --- | --- | --- | --- | --- | --- | --- | --- | --- | --- |
| ICEBERG (Goldman'24 [46]) | 0.203 | 0.383 | 0.492 | 0.565 | 0.617 | 0.658 | 0.693 | 0.722 | 0.745 | 0.765 |
| ICEBERG (Ours) |  |  |  |  |  |  |  |  |  |  |
| + collision energy | 0.214 | 0.411 | 0.537 | 0.610 | 0.662 | 0.704 | 0.739 | 0.767 | 0.791 | 0.809 |
| + entropy loss | 0.288 | 0.483 | 0.593 | 0.669 | 0.718 | 0.756 | 0.784 | 0.808 | 0.827 | 0.844 |
| + contrastive finetune | 0.294 | 0.498 | 0.611 | 0.680 | 0.729 | 0.757 | 0.785 | 0.806 | 0.826 | 0.842 |
| + $\times 2$ batch size | 0.307 | 0.503 | 0.612 | 0.677 | 0.728 | 0.762 | 0.791 | 0.809 | 0.826 | 0.844 |
| + charge migration | **0.335** | **0.543** | **0.658** | **0.730** | **0.774** | **0.811** | **0.842** | **0.865** | **0.880** | **0.892** |

We validate the source of accuracy improvements on the NIST'20 random split. Retrieval accuracy values are averaged among all positive and negative adduct types. The training random seed is set to 1 in this ablation study.

**Extended Data Table 2:**
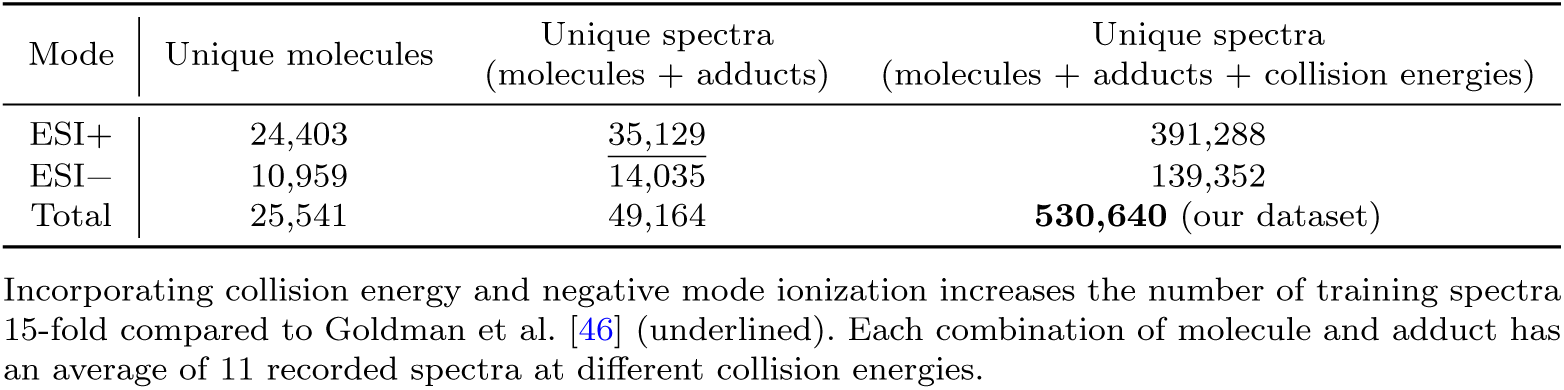
Statistics of the Orbitrap spectra from the NIST’20 library.

