## Supplementary Information for "Neural Spectral Prediction for Structure Elucidation with Tandem Mass Spectrometry"

**S4 Analytical standards** **26**

**S5 Elucidation of mental depression biomarkers** **29**

**S6 Elucidation of TB meningitis biomarkers and cost estimation of custom synthesis** **30**

**S7 Pesticide degradation experimentation** **31**

### Appendix S1 Discussion on rearrangements

Despite the fact that the ICEBERG model architecture does not consider rearrangements explicitly, we demonstrate that, interestingly, most known rearrangement mechanisms reported by Demarque et al. [53] are covered by ICEBERG's hydrogen shift predictions (Supplementary Figures S1 and S2). The structures are not necessarily correct, but the correct $m/z$ value is within ICEBERG's output range, making it capable of learning these rearrangement rules from data. Our newly introduced charge migration handling especially extends the coverage to charge migration rearrangements.

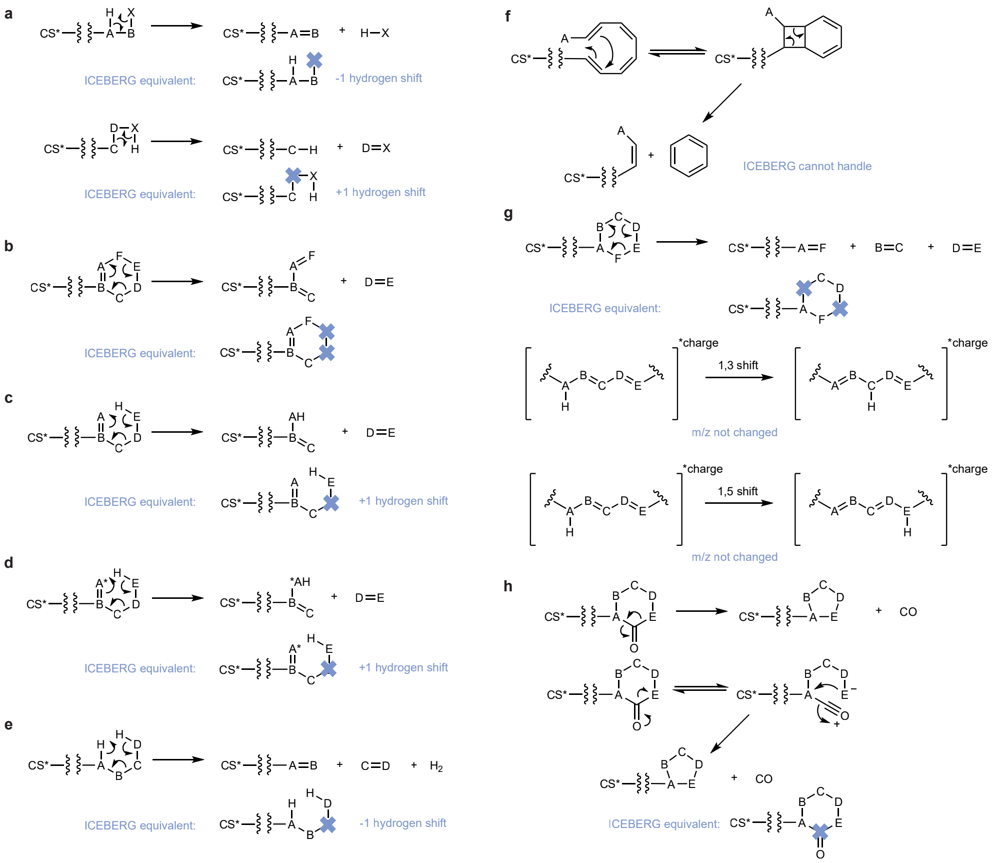

**Supplementary Fig. S1**: **ICEBERG is able to handle several known charge-retention rearrangements by predicting peaks with the correct $m/z$.** Among eight rearrangement mechanisms classified as charge retention fragmentation (CRF) by Demarque et al. [53], ICEBERG could predict fragments at the correct $m/z$ for seven. CS* denotes a charged site and could be positive or negative. The mechanisms are: **a**, Remote hydrogen rearrangements. **b**, Retro-Diels-Alder (RDA) reactions. **c**, Retro-ene reactions. **d**, Retro-heteroene reactions (*A is heteroatom). **e**, Charge remote fragmentation. **f**, Aromatic eliminations, which is the only mechanism ICEBERG cannot handle. **g**, Other pericyclic processes. **h**, Carbon monoxide eliminations from cyclic carbonyls compounds.

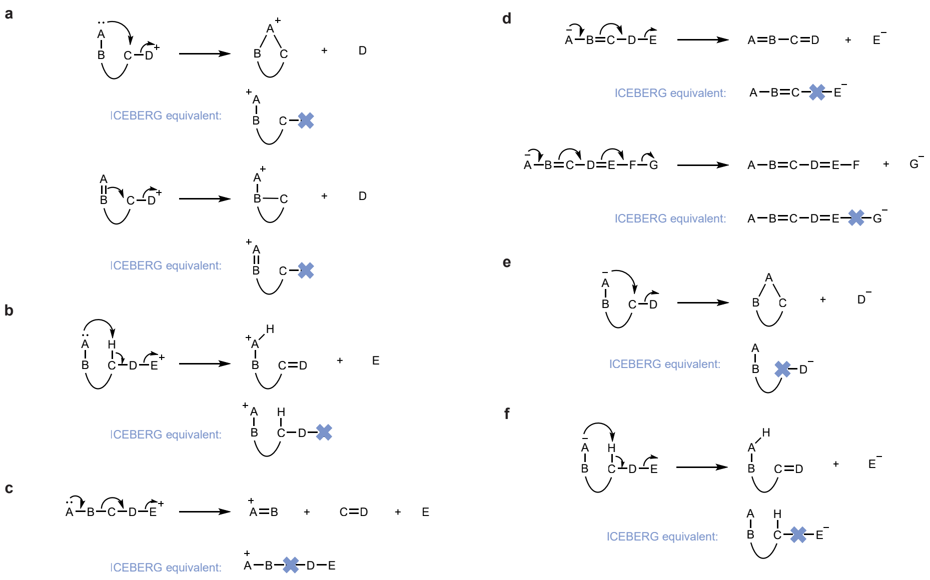

**Supplementary Fig. S2**: **ICEBERG handles known charge migration rearrangements by predicting peaks with the correct** $m/z$. Demarque et al. [53] further summarized six reaction mechanisms with charge migration fragmentation (CMF), where ICEBERG could predict fragments at the correct $m/z$ for all of them. **a**, Displacement reactions (positive ions). **b**, Inductive cleavages assisted by $\beta$-hydrogen removal. **c**, Grob-Wharton fragmentation. **d**, $\beta$-and $\epsilon$-eliminations. **e**, Displacement reactions (negative ions). **f**, Eliminations assisted by $\beta$-hydrogen removal.

### Appendix S2 Quantitative evaluation details

#### S2.1 MS/MS predictor baselines (on the NIST'20 benchmark)

We reimplement GrAFF-MS [27] and MassFormer [29] and retrain them, where the training and testing scripts are also available in our open-source code repository. CFM-ID supports $[M+H]^+$ and $[M-H]^-$ adducts. As a reproducibility note, we use a pretrained version of CFM-ID [25, 26], which may introduce potential data leakage; however, since CFM-ID still underperforms compared to other methods, this concern is less significant. Due to the potential data leakage and the unavailability of retraining, CFM-ID is excluded from the retrieval benchmark. Instead, we evaluate a learning-free fragmentation heuristic, MetFrag [23, 105] (version 2.6.5), with the default parameters on the same NIST'20 testing dataset. MetFrag does not have a structural prediction module, therefore, we only evaluate its retrieval accuracy. We assign the same PubChem candidate set (described in Supplementary Section S2.2) to MetFrag for fair comparison. FraGNNet [30] results are quoted directly from the original paper, as open-source code is not available at the time of writing. Moreover, since FraGNNet does not report retrieval accuracy under the scaffold split setting, Extended Data Fig. 2d omits FraGNNet from retrieval comparisons. ICEBERG (Goldman'24) refers to the original version of ICEBERG in Goldman et al. [46], without the technical improvements introduced in this paper. GrAFF-MS and MassFormer are trained on the same dataset described in Methods. FraGNNet is trained exclusively on $[M+H]^+$ spectra according to its publication, while ICEBERG (Goldman'24) was trained only on positive mode spectra, as reported in the original work.

#### S2.2 Retrieval benchmark setup (on the NIST'20 benchmark)

The retrieval benchmark is designed as a realistic *in silico* evaluation (i.e., retrospective analysis) of MS/MS predictors in real-world applications. In this benchmark, a spectrum predictor such as ICEBERG ranks candidate structures based on the similarity between predicted and experimental spectra. We evaluate whether the correct molecule is ranked within the top-$k$ predicted candidates and report detailed quantitative results (Supplementary Tables S2, S4, S6 and S7). Since the molecular formula of an unknown compound can often be inferred independently using tools like MIST-CF [102], SIRIUS [59], or Buddy [101], we restrict candidate structures to those with the same formula. Following Goldman et al. [46], we construct the candidate set using up to 49 isomers from PubChem. These are selected by ranking all PubChem structures with the same formula according to their Tanimoto similarity [106] to the ground-truth structure. The selected isomers, along with the ground truth, form a pool of up to 50 candidates. Such a design prioritizes decoy structures that are most structurally similar in the Tanimoto space, under the assumption that structurally similar molecules are more likely to have similar fragmentation patterns; therefore, the most challenging 49 structures are selected as candidates.

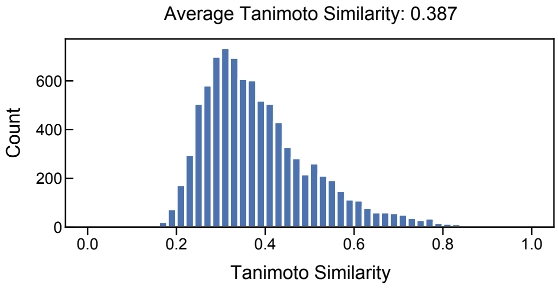

**Supplementary Fig. S3**: Tanimoto similarity distribution between the MSnLib test set and the closest NIST structure.

#### S2.3 Evaluation with SIRIUS on MSnLib

We evaluate the structural elucidation accuracy of ICEBERG with the off-the-shelf computational tool, SIRIUS (6.0.6) [37] (Fig. 2c). We focused on the most established workflow on SIRIUS: using CSI:FingerID [36] to predict the molecular fingerprint from MS/MS and use the fingerprint to rank candidate structures. We included two configurations: (1) the default configuration of SIRIUS where it only looks up a bio-relevant structures (from Biocyc, Blood Exposome, CHEBI, COCONUT, FooDB, GNPS, HMDB, HSDB, KEGG, KNApSAcK, LOTUS, LipidMaps, Maconda, MeSH, MiMeDB, NORMAN, Plantcyc, SuperNatural, TeroMOL, YMDB, and PubChem subsets of bio and metabolites, drug, food, and safety and toxic); and (2) a fair comparison configuration where SIRIUS looks up all structures from PubChem (a superset of all), same as the ICEBERG elucidation workflow. We also evaluated the *de novo* generation workflow, MSNovelist [38].

Since SIRIUS is a commercial software that does not support easy retraining, it is hard to find a fair testing split in NIST'20 without the concern of data leakage. We utilized a recently available resource, the MSnLib collected by Brungs et al. [50], containing 8,519 MS2 spectra for 7,973 unique structures that are not recorded in NIST or GNPS. We also exclude spectra with rare adduct types not supported by ICEBERG (10), where SIRIUS failed to make predictions (176), and where SIRIUS predicted the wrong formula (128). Although both SIRIUS and ICEBERG use PubChem candidates, their coverage is slightly different: there are 42 spectra whose structures do not exist in the PubChem cached by SIRIUS 6.0.6, and there are 55 spectra not included in our cached PubChem. We keep those entries in testing to reflect the real-world challenge, while giving SIRIUS a slight advantage over ICEBERG. We use the ICEBERG model trained on NIST'20 random split with random seed = 1, same as the model reported in Fig. 2a. To understand the distribution difference between MSnLib and NIST, we measure the Tanimoto similarity (of Morgan fingerprint, 2048 bits, radius 3) between the MSnLib structures and their closest strucutres in NIST, as reported in Supplementary Fig. S3. The detailed accuracy statistics are reported in Supplementary Table S10.

#### S2.4 Evaluation of top-1 retrieval accuracy

We bin the testing samples according to molecular mass and the average number of peaks across different collision energies (Fig. 2d). For each bin, the top-1 retrieval accuracy is calculated and visualized. Circle size reflects the number of testing samples in the bin, while color indicates top-1 retrieval accuracy. Marginal distributions of molecular mass and peak count are shown alongside. Results are reported for the random split, $[M+H]^+$ adduct, and training random seed = 1. The detailed statistics for each bin are reported in Supplementary Tables S11 and S12.

#### S2.5 Evaluation of entropy similarity and molecular mass

We illustrate the correlation between the predicted entropy similarity (to the ground-truth structure) and molecular mass (Fig. 2e). Darker hexagons indicate higher sample density within the bin. Marginal distributions are provided for both axes. Results correspond to the random split, $[M+H]^+$ adduct, and training random seed = 1. Since Fig. 2e is a hexagon plot where its detailed statistics are hard to show as a table, the code is available to recreate the same plot in our GitHub repository.

#### S2.6 Evaluation on MassSpecGym dataset

We utilize the open-source dataset collected by MassSpecGym [43] to train and benchmark a fully open-sourced version of ICEBERG (Fig. 2b). We adopted the data split developed by MassSpecGym authors, where all testing molecules have at least an MCES (Maximum Common Edge Subgraph) edit distance of 10 with any training molecules. Since ICEBERG requires collision energy as input and not all spectra in MassSpecGym have such an annotation, we only included spectra with collision energy labels for training and testing, which is in line with the setting used by FraGNNet [30] and the MassSpecGym "spectrum simulation challenge". We benchmarked cosine similarity, entropy similarity (also known as Jensen-Shannon similarity), and top-$k$ retrieval accuracies on the testing dataset. Results are reported with 99.9% confidence intervals upon bootstrapping, 20,000 resamples, following the MassSpecGym benchmark. The detailed similarities and accuracies are reported in Supplementary Table S13.

#### S2.7 Evaluation of accuracy per compound class

We calculate the top-$k$ retrieval accuracy for random split, seed = 1 (Extended Data Fig. 3a). Compound classes are predicted by NPClassifier [104], and then ordered by descending top-1 accuracy. The detailed per-class retrieval accuracy statistics are reported in Supplementary Table S14.

#### S2.8 Evaluation of accuracy per adduct type

We calculate the average top-$k$ retrieval accuracy across three random seeds (Extended Data Fig. 3b). Adduct types are ordered by descending top-1 accuracy, and the number of training samples for each adduct type is shown on the second y-axis. The detailed per-adduct retrieval accuracy statistics are reported in Supplementary Tables S15 and S16.

#### S2.9 Extended quantitative results on NIST'20

We report extended evaluation results for the best reproducibility of Fig. 2a and Extended Data Fig. 2d, including detailed cosine similarity, entropy similarity, and peak coverage for $[M+H]^+$ adduct in Supplementary Table S1. Detailed top-$k$ retrieval accuracies are reported in Supplementary Table S2.

**Supplementary Table S1**: Detailed evaluation results on spectral prediction accuracy for $[M+H]^+$ adduct on NIST'20 dataset. ±: 95% CI.

| Method | **[M+H]⁺, Random split** | | |
|---|---|---|---|
| | Cosine sim. (↑) | Entropy sim. (↑) | Coverage (↑) |
| CFM-ID [26] | 0.470 ± 0.000 | 0.427 ± 0.000 | 0.261 ± 0.000 |
| GrAFF-MS [27] | 0.565 ± 0.001 | 0.529 ± 0.001 | 0.740 ± 0.001 |
| MassFormer [29] | 0.632 ± 0.001 | 0.605 ± 0.000 | 0.787 ± 0.001 |
| FraGNNet [30] | 0.717 ± 0.001 | - | - |
| ICEBERG (Goldman'24 [46]) | 0.754 ± 0.002 | 0.700 ± 0.001 | 0.809 ± 0.001 |
| **ICEBERG (Ours)** | **0.794** ± **0.002** | **0.763** ± **0.001** | **0.856** ± **0.000** |

| Method | **[M+H]⁺, Scaffold split** | | |
|---|---|---|---|
| | Cosine sim. (↑) | Entropy sim. (↑) | Coverage (↑) |
| CFM-ID [26] | 0.450 ± 0.000 | 0.401 ± 0.000 | 0.228 ± 0.000 |
| GrAFF-MS [27] | 0.470 ± 0.001 | 0.448 ± 0.001 | 0.706 ± 0.001 |
| MassFormer [29] | 0.526 ± 0.002 | 0.512 ± 0.002 | 0.742 ± 0.002 |
| FraGNNet [30] | 0.654 ± 0.003 | - | - |
| ICEBERG (Goldman'24 [46]) | 0.701 ± 0.001 | 0.641 ± 0.004 | 0.790 ± 0.001 |
| **ICEBERG (Ours)** | **0.735** ± **0.000** | **0.708** ± **0.000** | **0.847** ± **0.000** |

As reported in this work, ICEBERG-predicted mass spectra are more similar to the experimental spectra than peer methods. As a general trend, entropy similarities tend to score worse compared to cosine similarities, while switching to the other metric does not affect the relative ranking of methods considered. ICEBERG-predicted peaks also provide higher coverage (i.e., recall) of experiment peaks. FraGNNet [30] is not open-sourced by the time this paper is written, therefore, its entropy similarity and coverage results are not available. "Coverage" denotes the recall in terms of the number of peaks annotated by the model divided by the number of experimental peaks. Results are reported with 95% CI on 3 random seeds.

We also report detailed results on other supported adduct types beyond $[M+H]^+$ (Extended Data Fig. 2e-j). Similarly, we report detailed evaluation results on positive mode in Supplementary Tables S3 and S4. We report detailed results on all adduct types in Supplementary Tables S5 and S6. For a fair comparison with MetFrag, we report the detailed retrieval accuracy with all adduct types MetFrag supports in Supplementary Table S7. We demonstrate that ICEBERG not only reaches state-of-the-art accuracy on $[M+H]^+$ but also outperforms baselines on other adduct types. There are fewer training data points for other adduct types, therefore, it is not too surprising

**Supplementary Table S2**: Detailed retrieval accuracy for $[M+H]^+$ adduct on NIST'20 dataset. ±: 95% CI.

| | | | $[M+H]^+$, **Random split** | | |
|---|---|---|---|---|---|
| Top-$k$ accuracy | Top-1 | Top-2 | Top-3 | Top-4 | Top-5 |
| MetFrag [105] | 0.107 ± 0.000 | 0.193 ± 0.000 | 0.267 ± 0.000 | 0.317 ± 0.000 | 0.375 ± 0.000 |
| GrAFF-MS [27] | 0.211 ± 0.004 | 0.365 ± 0.009 | 0.472 ± 0.015 | 0.551 ± 0.013 | 0.608 ± 0.005 |
| MassFormer [29] | 0.252 ± 0.001 | 0.422 ± 0.002 | 0.539 ± 0.005 | 0.617 ± 0.007 | 0.675 ± 0.004 |
| FraGNNet [30] | 0.238 | - | 0.504 | - | 0.652 |
| ICEBERG (Goldman'24 [46]) | 0.251 ± 0.016 | 0.454 ± 0.004 | 0.576 ± 0.006 | 0.654 ± 0.004 | 0.711 ± 0.007 |
| **ICEBERG (Ours)** | **0.400 ± 0.008** | **0.611 ± 0.013** | **0.721 ± 0.010** | **0.790 ± 0.007** | **0.829 ± 0.006** |
| Top-$k$ accuracy | Top-6 | Top-7 | Top-8 | Top-9 | Top-10 |
| MetFrag [105] | 0.414 ± 0.000 | 0.457 ± 0.000 | 0.497 ± 0.000 | 0.537 ± 0.000 | 0.567 ± 0.000 |
| GrAFF-MS [27] | 0.655 ± 0.006 | 0.690 ± 0.004 | 0.724 ± 0.008 | 0.754 ± 0.010 | 0.776 ± 0.009 |
| MassFormer [29] | 0.724 ± 0.005 | 0.761 ± 0.008 | 0.794 ± 0.010 | 0.822 ± 0.005 | 0.843 ± 0.006 |
| FraGNNet [30] | - | - | - | - | 0.831 |
| ICEBERG (Goldman'24 [46]) | 0.753 ± 0.007 | 0.783 ± 0.005 | 0.810 ± 0.001 | 0.833 ± 0.007 | 0.850 ± 0.009 |
| **ICEBERG (Ours)** | **0.859 ± 0.005** | **0.881 ± 0.006** | **0.900 ± 0.009** | **0.913 ± 0.008** | **0.923 ± 0.007** |

| | | | $[M+H]^+$, **Scaffold split** | | |
|---|---|---|---|---|---|
| Top-$k$ accuracy | Top-1 | Top-2 | Top-3 | Top-4 | Top-5 |
| MetFrag [105] | 0.127 ± 0.000 | 0.225 ± 0.000 | 0.302 ± 0.000 | 0.363 ± 0.000 | 0.420 ± 0.000 |
| GrAFF-MS [27] | 0.143 ± 0.004 | 0.270 ± 0.002 | 0.368 ± 0.008 | 0.450 ± 0.006 | 0.518 ± 0.002 |
| MassFormer [29] | 0.184 ± 0.008 | 0.328 ± 0.005 | 0.431 ± 0.001 | 0.512 ± 0.005 | 0.575 ± 0.004 |
| ICEBERG (Goldman'24 [46]) | 0.229 ± 0.004 | 0.415 ± 0.006 | 0.535 ± 0.008 | 0.617 ± 0.009 | 0.675 ± 0.004 |
| **ICEBERG (Ours)** | **0.335 ± 0.005** | **0.543 ± 0.004** | **0.669 ± 0.006** | **0.745 ± 0.002** | **0.799 ± 0.004** |
| Top-$k$ accuracy | Top-6 | Top-7 | Top-8 | Top-9 | Top-10 |
| MetFrag [105] | 0.471 ± 0.000 | 0.510 ± 0.000 | 0.548 ± 0.000 | 0.582 ± 0.000 | 0.612 ± 0.000 |
| GrAFF-MS [27] | 0.569 ± 0.002 | 0.614 ± 0.003 | 0.650 ± 0.000 | 0.687 ± 0.002 | 0.716 ± 0.002 |
| MassFormer [29] | 0.629 ± 0.004 | 0.674 ± 0.006 | 0.712 ± 0.004 | 0.749 ± 0.004 | 0.775 ± 0.004 |
| ICEBERG (Goldman'24 [46]) | 0.721 ± 0.001 | 0.756 ± 0.003 | 0.787 ± 0.001 | 0.812 ± 0.002 | 0.834 ± 0.002 |
| **ICEBERG (Ours)** | **0.838 ± 0.006** | **0.867 ± 0.005** | **0.890 ± 0.006** | **0.907 ± 0.009** | **0.920 ± 0.009** |

The ICEBERG model presented in this work outperforms all peer methods in retrieval accuracy, improving the top-1 accuracy significantly from 25.1% (95% CI 23.5–26.7%) to 40.0% (95% CI 39.2–40.8%). FraGNNet results are quoted from their manuscript [30]. Results are reported with 95% CI on 3 random seeds.

to see a slight performance drop for all methods when more (relatively) uncommon adduct types are involved. We report in Supplementary Table S8 an ablation study on proton and non-proton adducts to highlight the impact of charge migration on model performance. In Supplementary Table S9, we compare two design choices of using positional encoding or normalized collision energy to incorporate the collision energy. The performance ranking of peer methods is quite consistent among different adduct types and different evaluation metrics within the scope of this paper.

**Supplementary Table S3**: Detailed evaluation results on spectral prediction accuracy for positive mode adducts on NIST'20 dataset. ±: 95% CI from three random seeds.

| Method | **Positive mode, Random split** | | |
|---|---|---|---|
| | Cosine sim. (↑) | Entropy sim. (↑) | Coverage (↑) |
| GrAFF-MS [27] | 0.578 ± 0.001 | 0.533 ± 0.001 | 0.724 ± 0.000 |
| MassFormer [29] | 0.640 ± 0.001 | 0.605 ± 0.000 | 0.776 ± 0.001 |
| ICEBERG (Goldman'24 [46]) | 0.727 ± 0.002 | 0.664 ± 0.000 | 0.754 ± 0.002 |
| **ICEBERG (Ours)** | **0.782** ± **0.002** | **0.749** ± **0.001** | **0.837** ± **0.000** |

| Method | **Positive mode, Scaffold split** | | |
|---|---|---|---|
| | Cosine sim. (↑) | Entropy sim. (↑) | Coverage (↑) |
| GrAFF-MS [27] | 0.477 ± 0.001 | 0.452 ± 0.001 | 0.701 ± 0.001 |
| MassFormer [29] | 0.532 ± 0.002 | 0.515 ± 0.002 | 0.740 ± 0.001 |
| ICEBERG (Goldman'24 [46]) | 0.698 ± 0.001 | 0.633 ± 0.004 | 0.770 ± 0.001 |
| **ICEBERG (Ours)** | **0.734** ± **0.000** | **0.707** ± **0.000** | **0.842** ± **0.000** |

**Supplementary Table S4**: Detailed retrieval accuracy for positive mode adducts on NIST'20 dataset. ±: 95% CI from three random seeds.

| | **Positive mode, Random split** | | | | |
|---|---|---|---|---|---|
| Top-$k$ accuracy | Top-1 | Top-2 | Top-3 | Top-4 | Top-5 |
| GrAFF-MS [27] | 0.173 ± 0.004 | 0.316 ± 0.003 | 0.417 ± 0.007 | 0.490 ± 0.006 | 0.545 ± 0.000 |
| MassFormer [29] | 0.219 ± 0.002 | 0.369 ± 0.004 | 0.476 ± 0.005 | 0.551 ± 0.006 | 0.607 ± 0.005 |
| ICEBERG (Goldman'24 [46]) | 0.189 ± 0.012 | 0.375 ± 0.005 | 0.489 ± 0.007 | 0.567 ± 0.005 | 0.623 ± 0.004 |
| **ICEBERG (Ours)** | **0.340 ± 0.004** | **0.550 ± 0.011** | **0.661 ± 0.005** | **0.735 ± 0.006** | **0.777 ± 0.006** |
| | **Positive mode, Random split** | | | | |
| Top-$k$ accuracy | Top-6 | Top-7 | Top-8 | Top-9 | Top-10 |
| GrAFF-MS [27] | 0.591 ± 0.003 | 0.625 ± 0.002 | 0.660 ± 0.004 | 0.690 ± 0.005 | 0.714 ± 0.006 |
| MassFormer [29] | 0.654 ± 0.004 | 0.693 ± 0.004 | 0.725 ± 0.004 | 0.755 ± 0.001 | 0.775 ± 0.004 |
| ICEBERG (Goldman'24 [46]) | 0.666 ± 0.002 | 0.698 ± 0.002 | 0.725 ± 0.003 | 0.749 ± 0.003 | 0.770 ± 0.002 |
| **ICEBERG (Ours)** | **0.810 ± 0.006** | **0.834 ± 0.008** | **0.855 ± 0.011** | **0.871 ± 0.008** | **0.885 ± 0.008** |

| | **Positive mode, Scaffold split** | | | | |
|---|---|---|---|---|---|
| Top-$k$ accuracy | Top-1 | Top-2 | Top-3 | Top-4 | Top-5 |
| GrAFF-MS [27] | 0.138 ± 0.003 | 0.262 ± 0.005 | 0.358 ± 0.008 | 0.437 ± 0.006 | 0.502 ± 0.003 |
| MassFormer [29] | 0.179 ± 0.007 | 0.319 ± 0.004 | 0.421 ± 0.003 | 0.499 ± 0.003 | 0.562 ± 0.003 |
| ICEBERG (Goldman'24 [46]) | 0.210 ± 0.004 | 0.397 ± 0.006 | 0.518 ± 0.006 | 0.599 ± 0.007 | 0.657 ± 0.002 |
| **ICEBERG (Ours)** | **0.321 ± 0.006** | **0.525 ± 0.003** | **0.650 ± 0.007** | **0.724 ± 0.005** | **0.778 ± 0.005** |
| Top-$k$ accuracy | Top-6 | Top-7 | Top-8 | Top-9 | Top-10 |
| GrAFF-MS [27] | 0.553 ± 0.002 | 0.598 ± 0.003 | 0.633 ± 0.001 | 0.670 ± 0.002 | 0.699 ± 0.001 |
| MassFormer [29] | 0.615 ± 0.004 | 0.660 ± 0.007 | 0.698 ± 0.006 | 0.734 ± 0.005 | 0.761 ± 0.006 |
| ICEBERG (Goldman'24 [46]) | 0.703 ± 0.001 | 0.738 ± 0.003 | 0.769 ± 0.001 | 0.794 ± 0.003 | 0.817 ± 0.003 |
| **ICEBERG (Ours)** | **0.818 ± 0.005** | **0.848 ± 0.005** | **0.871 ± 0.004** | **0.889 ± 0.008** | **0.903 ± 0.008** |

**Supplementary Table S5**: Detailed evaluation results on spectral prediction accuracy for both positive and negative mode adducts on NIST'20 dataset. ±: 95% CI from three random seeds.

| Method | **Positive+negative mode, Random split** | | |
|---|---|---|---|
| | Cosine sim. (↑) | Entropy sim. (↑) | Coverage (↑) |
| GrAFF-MS [27] | 0.556 ± 0.001 | 0.512 ± 0.001 | 0.710 ± 0.000 |
| MassFormer [29] | 0.656 ± 0.001 | 0.610 ± 0.000 | 0.782 ± 0.001 |
| **ICEBERG (Ours)** | **0.773 ± 0.002** | **0.740 ± 0.002** | **0.834 ± 0.001** |
| Method | **Positive+negative mode, Scaffold split** | | |
| | Cosine sim. (↑) | Entropy sim. (↑) | Coverage (↑) |
| GrAFF-MS [27] | 0.474 ± 0.001 | 0.448 ± 0.001 | 0.692 ± 0.001 |
| MassFormer [29] | 0.536 ± 0.001 | 0.516 ± 0.002 | 0.738 ± 0.002 |
| **ICEBERG (Ours)** | **0.733 ± 0.000** | **0.706 ± 0.000** | **0.837 ± 0.000** |

**Supplementary Table S6**: Detailed retrieval accuracy for both positive and negative mode adducts on NIST'20 dataset. ±: 95% CI from three random seeds.

| | **Positive+negative mode, Random split** | | | | |
|---|---|---|---|---|---|
| Top-$k$ accuracy | Top-1 | Top-2 | Top-3 | Top-4 | Top-5 |
| GrAFF-MS [27] | 0.165 ± 0.003 | 0.305 ± 0.003 | 0.406 ± 0.004 | 0.479 ± 0.005 | 0.535 ± 0.001 |
| MassFormer [29] | 0.212 ± 0.004 | 0.360 ± 0.005 | 0.465 ± 0.005 | 0.538 ± 0.004 | 0.595 ± 0.006 |
| **ICEBERG (Ours)** | **0.335 ± 0.000** | **0.549 ± 0.009** | **0.662 ± 0.004** | **0.733 ± 0.005** | **0.777 ± 0.004** |
| Top-$k$ accuracy | Top-6 | Top-7 | Top-8 | Top-9 | Top-10 |
| GrAFF-MS [27] | 0.582 ± 0.004 | 0.618 ± 0.001 | 0.653 ± 0.004 | 0.683 ± 0.007 | 0.708 ± 0.007 |
| MassFormer [29] | 0.642 ± 0.005 | 0.682 ± 0.004 | 0.715 ± 0.003 | 0.744 ± 0.001 | 0.765 ± 0.003 |
| **ICEBERG (Ours)** | **0.811 ± 0.002** | **0.838 ± 0.005** | **0.858 ± 0.008** | **0.875 ± 0.006** | **0.889 ± 0.005** |

| | **Positive+negative mode, Scaffold split** | | | | |
|---|---|---|---|---|---|
| Top-k accuracy | Top-1 | Top-2 | Top-3 | Top-4 | Top-5 |
| GrAFF-MS [27] | 0.140 ± 0.004 | 0.268 ± 0.005 | 0.364 ± 0.007 | 0.442 ± 0.005 | 0.506 ± 0.003 |
| MassFormer [29] | 0.179 ± 0.007 | 0.319 ± 0.004 | 0.420 ± 0.003 | 0.497 ± 0.002 | 0.559 ± 0.003 |
| **ICEBERG (Ours)** | **0.316 ± 0.007** | **0.524 ± 0.003** | **0.648 ± 0.005** | **0.720 ± 0.004** | **0.775 ± 0.005** |
| Top-k accuracy | Top-6 | Top-7 | Top-8 | Top-9 | Top-10 |
| GrAFF-MS [27] | 0.556 ± 0.001 | 0.599 ± 0.004 | 0.634 ± 0.003 | 0.670 ± 0.003 | 0.698 ± 0.001 |
| MassFormer [29] | 0.611 ± 0.003 | 0.656 ± 0.006 | 0.694 ± 0.005 | 0.730 ± 0.005 | 0.757 ± 0.006 |
| **ICEBERG (Ours)** | **0.814 ± 0.003** | **0.843 ± 0.004** | **0.866 ± 0.004** | **0.885 ± 0.007** | **0.899 ± 0.008** |

**Supplementary Table S7**: Detailed retrieval accuracy for MetFrag-supported adducts on NIST'20 dataset. ±: 95% CI from three random seeds.

| | **Positive+negative mode, Random split** | | | | |
|---|---|---|---|---|---|
| Top-$k$ accuracy | Top-1 | Top-2 | Top-3 | Top-4 | Top-5 |
| MetFrag [105] | 0.147 ± 0.000 | 0.237 ± 0.000 | 0.308 ± 0.000 | 0.370 ± 0.000 | 0.428 ± 0.000 |
| GrAFF-MS [27] | 0.178 ± 0.003 | 0.320 ± 0.004 | 0.423 ± 0.003 | 0.498 ± 0.004 | 0.556 ± 0.001 |
| MassFormer [29] | 0.222 ± 0.002 | 0.377 ± 0.004 | 0.486 ± 0.007 | 0.559 ± 0.002 | 0.617 ± 0.004 |
| **ICEBERG (Ours)** | **0.365 ± 0.001** | **0.582 ± 0.010** | **0.696 ± 0.005** | **0.765 ± 0.004** | **0.806 ± 0.004** |
| Top-$k$ accuracy | Top-6 | Top-7 | Top-8 | Top-9 | Top-10 |
| MetFrag [105] | 0.469 ± 0.000 | 0.514 ± 0.000 | 0.552 ± 0.000 | 0.588 ± 0.000 | 0.618 ± 0.000 |
| GrAFF-MS [27] | 0.603 ± 0.001 | 0.639 ± 0.002 | 0.673 ± 0.005 | 0.703 ± 0.007 | 0.727 ± 0.007 |
| MassFormer [29] | 0.665 ± 0.004 | 0.704 ± 0.005 | 0.736 ± 0.005 | 0.764 ± 0.002 | 0.785 ± 0.006 |
| **ICEBERG (Ours)** | **0.838 ± 0.003** | **0.864 ± 0.003** | **0.882 ± 0.007** | **0.897 ± 0.006** | **0.909 ± 0.005** |
| | **Positive+negative mode, Scaffold split** | | | | |
| Top-k accuracy | Top-1 | Top-2 | Top-3 | Top-4 | Top-5 |
| MetFrag [105] | 0.150 ± 0.000 | 0.247 ± 0.000 | 0.325 ± 0.000 | 0.386 ± 0.000 | 0.441 ± 0.000 |
| GrAFF-MS [27] | 0.144 ± 0.004 | 0.272 ± 0.004 | 0.369 ± 0.007 | 0.448 ± 0.005 | 0.512 ± 0.002 |
| MassFormer [29] | 0.181 ± 0.007 | 0.323 ± 0.004 | 0.424 ± 0.003 | 0.502 ± 0.002 | 0.565 ± 0.002 |
| **ICEBERG (Ours)** | **0.323 ± 0.007** | **0.534 ± 0.005** | **0.660 ± 0.005** | **0.732 ± 0.001** | **0.787 ± 0.003** |
| Top-k accuracy | Top-6 | Top-7 | Top-8 | Top-9 | Top-10 |
| MetFrag [105] | 0.489 ± 0.000 | 0.528 ± 0.000 | 0.564 ± 0.000 | 0.599 ± 0.000 | 0.628 ± 0.000 |
| GrAFF-MS [27] | 0.563 ± 0.002 | 0.607 ± 0.004 | 0.642 ± 0.002 | 0.678 ± 0.002 | 0.706 ± 0.000 |
| MassFormer [29] | 0.618 ± 0.003 | 0.662 ± 0.005 | 0.701 ± 0.005 | 0.737 ± 0.004 | 0.763 ± 0.004 |
| **ICEBERG (Ours)** | **0.826 ± 0.004** | **0.855 ± 0.004** | **0.878 ± 0.005** | **0.896 ± 0.009** | **0.909 ± 0.009** |

**Supplementary Table S8**: Comparison of retrieval accuracies for proton and non-proton adducts before and after incorporating charge migration (NIST'20, random split, random seed = 1).

| | Top-1 | Top-2 | Top-3 | Top-4 | Top-5 | Top-6 | Top-7 | Top-8 | Top-9 | Top-10 |
|---|---|---|---|---|---|---|---|---|---|---|
| **Without charge migration** | | | | | | | | | | |
| Proton Adducts | 0.374 | 0.586 | 0.698 | 0.764 | 0.811 | 0.843 | 0.871 | 0.886 | 0.900 | 0.912 |
| Non-proton Adducts | 0.180 | 0.348 | 0.452 | 0.515 | 0.574 | 0.610 | 0.641 | 0.665 | 0.689 | 0.715 |
| **With charge migration** | | | | | | | | | | |
| Proton Adducts | 0.384 | 0.591 | 0.706 | 0.774 | 0.814 | 0.849 | 0.879 | 0.899 | 0.914 | 0.924 |
| Non-proton Adducts | 0.243 | 0.454 | 0.568 | 0.647 | 0.699 | 0.741 | 0.774 | 0.800 | 0.817 | 0.833 |

Among the 4925 spectra in the NIST'20 testing set, 3210 of them are either $[M{+}H]^+$ or $[M{-}H]^-$ adducts. Incorporating charge migration only introduces a marginal improvement for them, because it is handled by hydrogen shift. Charge migration has a significant contribution to those non-proton adduct types, such as $[M{+}Na]^+$ and $[M{-}H_2O{+}H]^+$.

**Supplementary Table S9**: Ablation study of different ways of incorporating collision energy.

| Method | Top-1 | Top-5 | Top-10 |
|---|---|---|---|
| ICEBERG baseline (Goldman'24) | 0.203 | 0.617 | 0.765 |
| Normalized collision energy input | 0.194 | 0.643 | 0.798 |
| Positional encoding input | 0.214 | 0.662 | 0.809 |

**Supplementary Table S10**: Top-$k$ accuracies (%) of ICEBERG and SIRIUS on MSnLib subset excluding NIST and GNPS (Fig. 2c).

| Top-$k$ | SIRIUS (PubChem) | SIRIUS (Bio-only) | SIRIUS (De novo) | ICEBERG (Ours) |
|---|---|---|---|---|
| Top-1 | 15.69 | 31.27 | 2.86 | 38.00 |
| Top-2 | 23.17 | 37.56 | 4.11 | 53.76 |
| Top-3 | 28.14 | 40.94 | 4.77 | 62.57 |
| Top-4 | 31.60 | 43.09 | 5.18 | 67.53 |
| Top-5 | 34.18 | 44.50 | 5.62 | 71.46 |
| Top-6 | 36.10 | 45.70 | 5.89 | 73.98 |
| Top-7 | 38.47 | 46.67 | 6.17 | 76.07 |
| Top-8 | 39.88 | 47.42 | 6.41 | 77.73 |
| Top-9 | 41.17 | 48.12 | 6.70 | 79.16 |
| Top-10 | 42.45 | 48.89 | 6.86 | 80.29 |

**Supplementary Table S11**: Number of samples for each bin in the analysis of top-1 retrieval accuracy associated with molecular mass and number of peaks (i.e., size of markers in Fig. 2d), on NIST'20 testing dataset, random split.

| | Molecular mass range | | | | | | | | | | | | | | | | | |
|---|---|---|---|---|---|---|---|---|---|---|---|---|---|---|---|---|---|---|
| Average number of peaks | [82, 111) | [111, 139) | [139, 168) | [168, 197) | [197, 225) | [225, 254) | [254, 283) | [283, 311) | [311, 340) | [340, 369) | [369, 397) | [397, 426) | [426, 455) | [455, 484) | [484, 512) | [512, 541) | [541, 570) | [570, 598] |
| [1.0, 3.7) | 8 | 7 | 1 | 3 | 6 | 3 | 1 | 1 | 3 | 2 | 1 | 1 | 0 | 0 | 0 | 0 | 1 | 0 |
| [3.7, 6.4) | 5 | 18 | 22 | 19 | 10 | 7 | 3 | 6 | 5 | 1 | 3 | 1 | 3 | 0 | 1 | 1 | 0 | 0 |
| [6.4, 9.0) | 2 | 11 | 17 | 23 | 25 | 18 | 14 | 6 | 5 | 1 | 3 | 2 | 1 | 0 | 1 | 1 | 2 | 0 |
| [9.0, 11.7) | 2 | 13 | 18 | 22 | 24 | 24 | 18 | 15 | 6 | 4 | 5 | 1 | 0 | 1 | 1 | 0 | 0 | 0 |
| [11.7, 14.4) | 1 | 9 | 16 | 16 | 22 | 24 | 19 | 14 | 9 | 8 | 8 | 3 | 0 | 1 | 0 | 0 | 0 | 1 |
| [14.4, 17.1) | 0 | 2 | 13 | 22 | 23 | 24 | 21 | 13 | 6 | 12 | 6 | 0 | 4 | 4 | 1 | 1 | 0 | 0 |
| [17.1, 19.8) | 1 | 5 | 13 | 21 | 29 | 20 | 22 | 15 | 10 | 5 | 2 | 5 | 0 | 0 | 1 | 0 | 1 | 0 |
| [19.8, 22.4) | 1 | 9 | 13 | 19 | 26 | 23 | 22 | 15 | 13 | 4 | 11 | 2 | 2 | 3 | 2 | 1 | 1 | 0 |
| [22.4, 25.1) | 0 | 0 | 2 | 10 | 16 | 16 | 13 | 18 | 8 | 7 | 5 | 4 | 3 | 2 | 1 | 0 | 1 | 0 |
| [25.1, 27.8) | 0 | 1 | 7 | 20 | 18 | 15 | 22 | 24 | 13 | 11 | 8 | 4 | 0 | 1 | 2 | 0 | 2 | 0 |
| [27.8, 30.5) | 0 | 0 | 6 | 10 | 16 | 20 | 28 | 12 | 15 | 4 | 5 | 5 | 1 | 1 | 2 | 0 | 3 | 3 |
| [30.5, 33.1) | 0 | 2 | 5 | 8 | 21 | 21 | 16 | 15 | 13 | 8 | 7 | 4 | 4 | 1 | 4 | 3 | 2 | 0 |
| [33.1, 35.8) | 0 | 0 | 4 | 7 | 11 | 10 | 15 | 20 | 12 | 9 | 9 | 4 | 2 | 6 | 0 | 0 | 0 | 1 |
| [35.8, 38.5) | 0 | 0 | 2 | 3 | 7 | 16 | 23 | 13 | 12 | 13 | 8 | 2 | 6 | 6 | 1 | 1 | 1 | 5 |
| [38.5, 41.2) | 0 | 0 | 0 | 5 | 12 | 15 | 15 | 13 | 17 | 6 | 14 | 8 | 3 | 3 | 4 | 1 | 0 | 4 |
| [41.2, 43.9) | 0 | 0 | 1 | 0 | 7 | 9 | 16 | 16 | 14 | 11 | 8 | 5 | 4 | 3 | 3 | 0 | 5 | 1 |
| [43.9, 46.5) | 0 | 0 | 0 | 1 | 0 | 4 | 7 | 10 | 11 | 11 | 5 | 6 | 2 | 6 | 3 | 3 | 0 | 1 |
| [46.5, 49.2] | 0 | 0 | 0 | 0 | 0 | 0 | 2 | 3 | 3 | 3 | 1 | 1 | 0 | 2 | 1 | 0 | 0 | 0 |

**Supplementary Table S12**: Top-1 retrieval accuracy for each bin in the analysis associated with molecular mass and number of peaks (i.e., hue of markers in Fig. 2d). "-" denotes there is no testing sample in this bin.

| | Molecular mass range | | | | | | | | | | | | | | | | | |
|---|---|---|---|---|---|---|---|---|---|---|---|---|---|---|---|---|---|---|
| Average number of peaks | [82, 111) | [111, 139) | [139, 168) | [168, 197) | [197, 225) | [225, 254) | [254, 283) | [283, 311) | [311, 340) | [340, 369) | [369, 397) | [397, 426) | [426, 455) | [455, 484) | [484, 512) | [512, 541) | [541, 570) | [570, 598] |
| [1.0, 3.7) | 0.38 | 0.57 | 1.00 | 0.00 | 0.33 | 0.33 | 0.00 | 0.00 | 0.00 | 0.50 | 1.00 | 0.00 | - | - | - | - | 0.00 | - |
| [3.7, 6.4) | 0.40 | 0.33 | 0.64 | 0.58 | 0.60 | 0.29 | 0.33 | 0.50 | 0.20 | 0.00 | 0.33 | 0.00 | 0.33 | - | 0.00 | 0.00 | - | - |
| [6.4, 9.0) | 0.00 | 0.45 | 0.41 | 0.43 | 0.36 | 0.44 | 0.29 | 0.50 | 0.80 | 1.00 | 0.67 | 0.00 | 0.00 | - | 1.00 | 0.00 | 0.50 | - |
| [9.0, 11.7) | 1.00 | 0.54 | 0.50 | 0.45 | 0.42 | 0.46 | 0.50 | 0.20 | 0.83 | 0.00 | 0.60 | 0.00 | - | 0.00 | 0.00 | - | - | - |
| [11.7, 14.4) | 0.00 | 0.22 | 0.38 | 0.44 | 0.41 | 0.50 | 0.32 | 0.57 | 0.44 | 0.50 | 0.50 | 0.33 | - | 1.00 | - | - | - | 0.00 |
| [14.4, 17.1) | - | 0.50 | 0.46 | 0.32 | 0.52 | 0.33 | 0.48 | 0.38 | 0.33 | 0.58 | 1.00 | - | 0.25 | 0.25 | 1.00 | 0.00 | - | - |
| [17.1, 19.8) | 0.00 | 0.00 | 0.31 | 0.48 | 0.45 | 0.35 | 0.55 | 0.40 | 0.50 | 0.40 | 1.00 | 0.60 | - | - | 0.00 | - | 0.00 | - |
| [19.8, 22.4) | 1.00 | 0.22 | 0.31 | 0.37 | 0.23 | 0.57 | 0.36 | 0.47 | 0.31 | 0.25 | 0.36 | 0.50 | 0.00 | 0.67 | 0.50 | 0.00 | 1.00 | - |
| [22.4, 25.1) | - | - | 0.00 | 0.40 | 0.19 | 0.44 | 0.38 | 0.33 | 0.25 | 0.43 | 0.20 | 0.50 | 1.00 | 0.50 | 0.00 | - | 1.00 | - |
| [25.1, 27.8) | - | 0.00 | 0.43 | 0.25 | 0.67 | 0.53 | 0.36 | 0.62 | 0.38 | 0.36 | 0.38 | 0.25 | - | 1.00 | 0.00 | - | 0.00 | - |
| [27.8, 30.5) | - | - | 0.17 | 0.20 | 0.56 | 0.40 | 0.36 | 0.25 | 0.40 | 0.50 | 0.20 | 0.80 | 1.00 | 1.00 | 0.50 | - | 0.67 | 0.67 |
| [30.5, 33.1) | - | 0.00 | 0.40 | 0.25 | 0.33 | 0.57 | 0.38 | 0.40 | 0.31 | 0.25 | 0.14 | 0.50 | 0.75 | 0.00 | 0.25 | 0.67 | 0.00 | - |
| [33.1, 35.8) | - | - | 0.50 | 0.00 | 0.09 | 0.50 | 0.40 | 0.55 | 0.50 | 0.11 | 0.33 | 0.75 | 1.00 | 0.67 | - | - | - | 0.00 |
| [35.8, 38.5) | - | - | 1.00 | 0.33 | 0.43 | 0.44 | 0.48 | 0.38 | 0.67 | 0.38 | 0.50 | 0.00 | 0.33 | 0.33 | 0.00 | 0.00 | 0.00 | 0.00 |
| [38.5, 41.2) | - | - | - | 0.80 | 0.33 | 0.33 | 0.40 | 0.54 | 0.35 | 0.17 | 0.29 | 0.50 | 0.33 | 0.33 | 1.00 | 0.00 | - | 0.25 |
| [41.2, 43.9) | - | - | 0.00 | - | 0.57 | 0.44 | 0.50 | 0.31 | 0.29 | 0.27 | 0.38 | 0.40 | 0.25 | 0.67 | 0.33 | - | 0.20 | 1.00 |
| [43.9, 46.5) | - | - | - | 0.00 | - | 0.50 | 0.29 | 0.30 | 0.09 | 0.18 | 0.40 | 0.17 | 0.00 | 0.33 | 0.67 | 0.33 | - | 1.00 |
| [46.5, 49.2] | - | - | - | - | - | - | 0.00 | 0.33 | 0.00 | 1.00 | 1.00 | 1.00 | - | 0.50 | 1.00 | - | - | - |

**Supplementary Table S13**: Performance comparison of different models on the MassSpecGym dataset (formula challenge for the retrieval results). Jensen-Shannon Similarity is the same as Spectral Entropy Similarity [49]. Values in parentheses denote 99.9% confidence intervals.

| Model | Cosine Similarity ↑ | Jensen-Shannon Similarity ↑ | Hit Rate @ 1 ↑ | Hit Rate @ 5 ↑ | Hit Rate @ 20 ↑ |
|---|---|---|---|---|---|
| Precursor m/z | 0.15 (0.14–0.17) | 0.15 (0.14–0.16) | 2.09 (1.66–2.59) | 8.52 (7.65–9.53) | 22.65 (21.26–24.01) |
| NEIMS (FFN) [93] | 0.25 (0.24–0.26) | 0.24 (0.23–0.25) | 7.62 (6.77–8.54) | 22.70 (21.32–24.12) | 44.12 (42.51–45.75) |
| NEIMS (GNN) [107] | 0.19 (0.18–0.20) | 0.20 (0.19–0.20) | 3.63 (3.05–4.29) | 13.55 (12.46–14.68) | 33.77 (32.26–35.37) |
| FraGNNet [30] | 0.52 (0.51–0.53) | 0.47 (0.46–0.48) | 31.93 (30.40–33.50) | 63.20 (61.64–64.76) | 82.70 (81.45–83.93) |
| ICEBERG (Ours) | **0.58 (0.57–0.59)** | **0.48 (0.48–0.49)** | **45.95 (44.32–47.59)** | **75.43 (73.88–76.90)** | **91.69 (90.73–92.60)** |

**Supplementary Table S14**: Detailed retrieval accuracies for different compound classes (random split, seed = 1, NIST'20 dataset, Extended Data Fig. 3a).

| Compound class | Top-1 retrieval | Top-5 retrieval | Top-10 retrieval |
|---|---|---|---|
| Diarylheptanoids | 0.61 | 0.83 | 0.91 |
| Ornithine alkaloids | 0.50 | 0.88 | 1.00 |
| Small peptides | 0.49 | 0.87 | 0.94 |
| Fatty amides | 0.48 | 0.93 | 0.96 |
| Histidine alkaloids | 0.46 | 0.73 | 0.88 |
| Peptide alkaloids | 0.43 | 0.89 | 0.98 |
| Isoflavonoids | 0.40 | 0.63 | 0.90 |
| Lignans | 0.39 | 0.82 | 0.93 |
| Anthranilic acid alkaloids | 0.39 | 0.81 | 0.93 |
| Glycerophospholipids | 0.38 | 0.80 | 0.87 |
| Pseudoalkaloids | 0.36 | 0.79 | 0.90 |
| Nicotinic acid alkaloids | 0.36 | 0.79 | 0.92 |
| Polycyclic aromatic polyketides | 0.36 | 0.74 | 0.87 |
| Lysine alkaloids | 0.36 | 0.81 | 0.88 |
| Fatty esters | 0.35 | 0.79 | 0.91 |
| Tryptophan alkaloids | 0.35 | 0.80 | 0.91 |
| Cyclic polyketides | 0.34 | 0.76 | 0.86 |
| Fatty Acids and Conjugates | 0.34 | 0.72 | 0.84 |
| Nucleosides | 0.32 | 0.91 | 0.98 |
| Aromatic polyketides | 0.31 | 0.96 | 1.00 |
| Coumarins | 0.30 | 0.80 | 0.93 |
| Meroterpenoids | 0.30 | 0.73 | 0.86 |
| Eicosanoids | 0.28 | 0.80 | 0.89 |
| Monoterpenoids | 0.24 | 0.69 | 0.81 |
| Flavonoids | 0.24 | 0.80 | 0.93 |
| Phenolic acids (C6-C1) | 0.24 | 0.70 | 0.89 |
| Tyrosine alkaloids | 0.22 | 0.57 | 0.81 |
| Macrolides | 0.22 | 0.88 | 0.94 |
| Diterpenoids | 0.20 | 0.66 | 0.81 |
| Steroids | 0.19 | 0.62 | 0.79 |
| Sesquiterpenoids | 0.19 | 0.49 | 0.67 |
| Phenylpropanoids (C6-C3) | 0.18 | 0.76 | 0.92 |
| Xanthones | 0.17 | 0.78 | 0.87 |
| Triterpenoids | 0.14 | 0.53 | 0.69 |
| Saccharides | 0.12 | 0.62 | 0.77 |

**Supplementary Table S15**: Detailed retrieval accuracies and training sizes for different adduct types (random split, NIST'20 dataset, Extended Data Fig. 3b).

| Adduct type | Top-1 retrieval | Top-5 retrieval | Top-10 retrieval | # Training spectra |
|---|---|---|---|---|
| $[M+H]^{+}$ | 0.40 | 0.83 | 0.92 | 214331 |
| $[M-H]^{-}$ | 0.35 | 0.80 | 0.91 | 89875 |
| $[M+NH_3+H]^{+}$ | 0.33 | 0.81 | 0.92 | 6263 |
| $[M+K]^{+}$ | 0.31 | 0.76 | 0.93 | 622 |
| $[M+Na]^{+}$ | 0.27 | 0.73 | 0.84 | 25582 |
| $[M-H-H_2O]^{-}$ | 0.24 | 0.70 | 0.85 | 7720 |
| $[M-H_2O+H]^{+}$ | 0.24 | 0.69 | 0.82 | 58426 |
| $[M-H-CO_2]^{-}$ | 0.23 | 0.67 | 0.84 | 9504 |
| $[M+Cl]^{-}$ | 0.22 | 0.71 | 0.89 | 6032 |
| $[M-2H_2O+H]^{+}$ | 0.16 | 0.55 | 0.71 | 11396 |

**Supplementary Table S16**: Detailed retrieval accuracies and training sizes for different adduct types (scaffold split, NIST'20 dataset, Extended Data Fig. 3b).

| Adduct type | Top-1 retrieval | Top-5 retrieval | Top-10 retrieval | # Training spectra |
|---|---|---|---|---|
| $[M+H]^{+}$ | 0.33 | 0.80 | 0.92 | 185847 |
| $[M-H]^{-}$ | 0.27 | 0.74 | 0.85 | 94755 |
| $[M+Na]^{+}$ | 0.24 | 0.69 | 0.85 | 25460 |
| $[M-H_2O+H]^{+}$ | 0.20 | 0.55 | 0.70 | 64581 |
| $[M+NH_3+H]^{+}$ | 0.20 | 0.71 | 0.89 | 6626 |
| $[M-2H_2O+H]^{+}$ | 0.13 | 0.39 | 0.58 | 13344 |
| $[M+Cl]^{-}$ | 0.06 | 0.83 | 0.94 | 7238 |
| $[M+K]^{+}$ | 0.00 | 0.00 | 0.67 | 694 |
| $[M-H-H_2O]^{-}$ | 0.00 | 0.00 | 0.67 | 9679 |

$[M-H-CO_2]^{-}$ does not have any testing samples under the scaffold split on NIST'20 dataset.

### Appendix S3 Qualitative study details

#### S3.1 Fragmentation analysis with bond types and collision energies

To study the fragmentation pattern predicted by ICEBERG, we analyzed the learned bond-breaking patterns of different bond types across collision energies (Fig. 2f). This analysis is achieved by extracting the fragmentation SMARTS from the generated fragmentation graphs of the NIST'20 test set to help identify the type of broken bond. We propose to perform analysis based on the following score:

$$P_{\text{bond\_type}} = \frac{\frac{I_{\text{bond\_type}}}{N_{\text{bond\_type}}}}{\sum_{\forall i} \frac{I_{\text{bond\_type}_i}}{N_{\text{bond\_type}_i}}} \times 100\% \tag{C1}$$

where $I_{\text{bond\_type}}$ denotes the sum of intensities of all peaks contributed by breaking the current broken bond type (e.g., C-C bonds, C-O bonds), and $N_{\text{bond\_type}}$ denotes the number of peaks. The denominator sums over all bond types. The score is computed for each bond type at each collision energy.

#### S3.2 DFT calculations of inductive cleavage energy ($C$)

We visualize bond-breaking events for the true positives in ICEBERG's predictions—i.e., bonds that ICEBERG identifies as breakable, and whose resulting fragments are indeed observed experimentally at corresponding $m/z$ values. We annotate these bond-breaking events with DFT-calculated inductive cleavage energy (Extended Data Fig. 4b). We compute the theoretical bond cleavage energy in the gas phase as the energy difference between the sum of the fragment energies and the single-point energy of the precursor. For simplicity, we omit reaction coordinates, rearrangements, matrix effects, and charge/proton migration; therefore, $C$ does not perfectly model the physical fragmentation process but instead serves as a theoretical protocol to study ICEBERG's learned fragmentation patterns. The workflow begins by calculating the inductive collision energy ($C$) as the difference between the single-point electronic energy of the protonated precursor and the combined energies of the observed (cationic) and neutral fragments (for the $[M+H]^+$ adduct) (Supplementary Fig. S4).

Initial geometries were generated using default settings from Open Babel (100 cycles of force field cleanup and one permutation of fast rotor search, obabel version 3.1.1). All DFT calculations were performed using ORCA (6.0.0), OpenMPI (4.1.6), and GCC (12.2.20). Geometry optimization was carried out at the PBE0-D4/def2-TZVP level of theory with the RIJCOSX approximation, tight optimization convergence criteria, and frequency calculations to confirm the nature of stationary points. If a transition state was detected, the starting geometry was manually adjusted, and the geometry optimization was repeated. Subsequent single-point energy calculations were performed using PBE0-D4/def2-QZVP with the RIJCOSX approximation and tight SCF convergence criteria. The optimized geometries were checked for the presence of transition states, in which case geometry optimizations were rerun with slightly adjusted starting geometries. All raw input and output files, as well as results, are provided as supplementary data under the MassIVE repository (MSV000097986).

The outlier for **2** at 10 eV in Extended Data Fig. 4c might be a failure example of ICEBERG, but it might also be because inductive cleavage is not the only mechanism of bond-breaking in mass spectrometry. For example, all the charge retention fragmentation (CRF) mechanisms [53] are not well-characterized by inductive cleavage, as well as rearrangements.

#### S3.3 Visualization of ICEBERG predictions

The visualized MS/MS predictions for the compound **2** are at normalized collision energies (NCEs) 10%, 20%, 30%, 40%, and 50% (Fig. 2h). ICEBERG is designed to work with absolute eV values, where the transformation is done by Equation (1). Collision energies in eV are then rounded to the nearest integer.

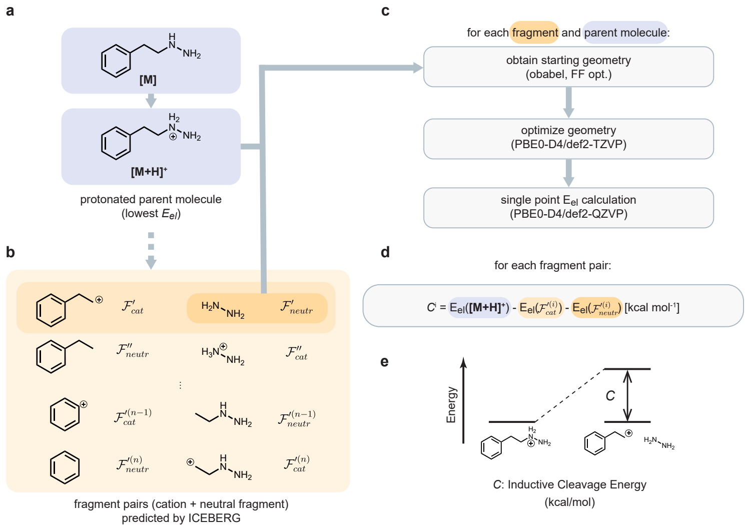

**Supplementary Fig. S4**: **Schematic overview of how an energy-based score helps to rationalize fragmentations by ICEBERG**. **a**, The protonated parent molecule $[M+H]^+$ of the neutral [M] is selected by choosing the protonation site with the lowest electronic energy in gas phase. **b**, For each molecule, a mass spectrum is generated using ICEBERG, and corresponding fragment pairs are labeled. Assuming heterolytic cleavage, each fragmentation event creates a neutral "protonated" fragment and a cationic fragment. **c**, The single point electronic energy is obtained by generating starting geometries using Open Babel, optimizing the geometry, and calculating the electronic energy with a fixed geometry. **d**, The inductive cleavage energy ($C$) score is computed by subtracting the energies of the complementary fragments from the energy of $[M+H]^+$. **e**, Illustration of $C$ with an energy diagram.

#### Appendix S4 Analytical standards

For the biomarker elucidation experiments, chemical standards were sourced from the following suppliers: hercynine (**1**) was obtained from 1Click Chemistry (Catalog No. CC169274, Lot No. 6PPVUR); LPC 19:0 (**9**) was purchased from Avanti Polar Lipids (Catalog No. 855776P); and GABA-Arg (**2**) was acquired from AnaSpec Inc. (Fremont, CA; Catalog No. SQ-ASPE-83805-2, 10 mg pack). For the novel pesticide degradant studies, carbendazim (**14**) was ordered from Chemspace, supplied by Enamine US (Chemspace ID: CSSB00011215494; IUPAC name: methyl N-(1H-1,3-benzodiazol-2-yl)carbamate; purity: 95%; quantity: 100 mg). Among them, GABA-Arg (**2**) is a non-standard compound made by a custom synthesis, whose NMR spectra are reported (Supplementary Figures S5 and S6).

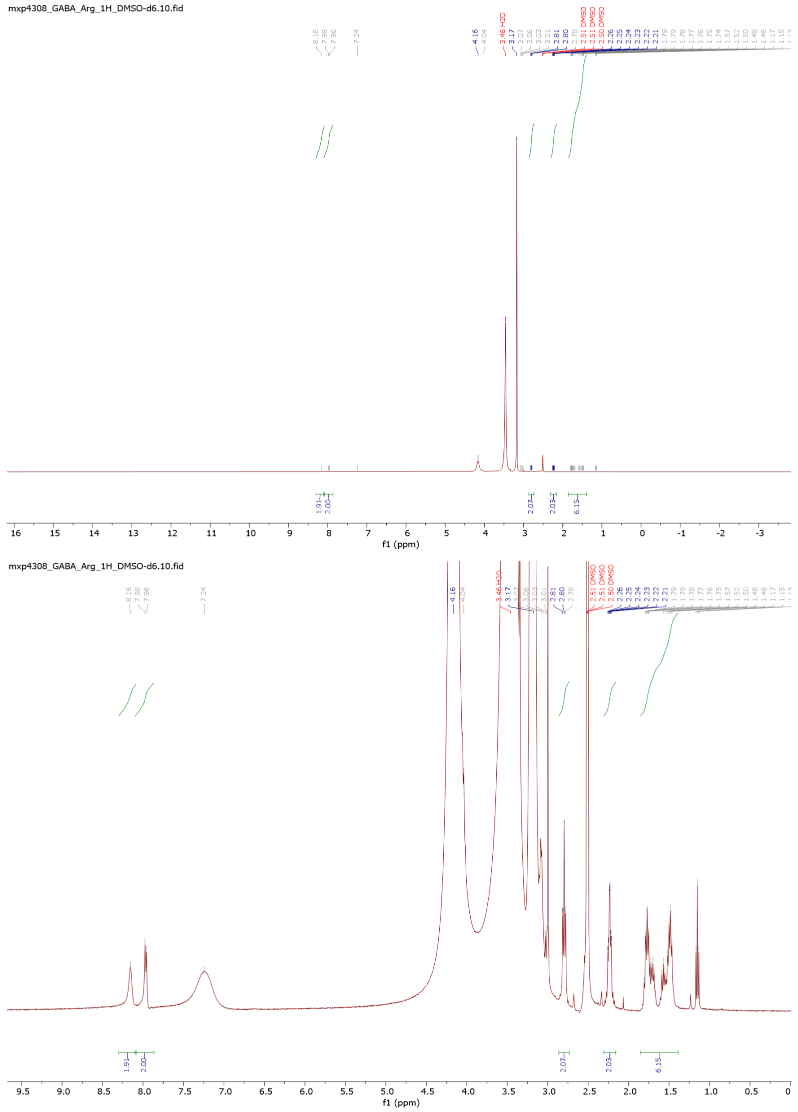

**Supplementary Fig. S5**: Original and zoomed-in $^{1}H$ spectra of **2** (400 MHz, DMSO-$d_6$): $\delta$ 8.16 (br s, 2H), 7.96 (br s, 2H), 2.8 (t, 3H), 2.26-2.21(t, 2H), 1.81-1.46 (m, 6H), 2.79-2.74 (m, 2H), 2.04-1.97 (m, 6H), 1.52-1.39 (m, 6H).

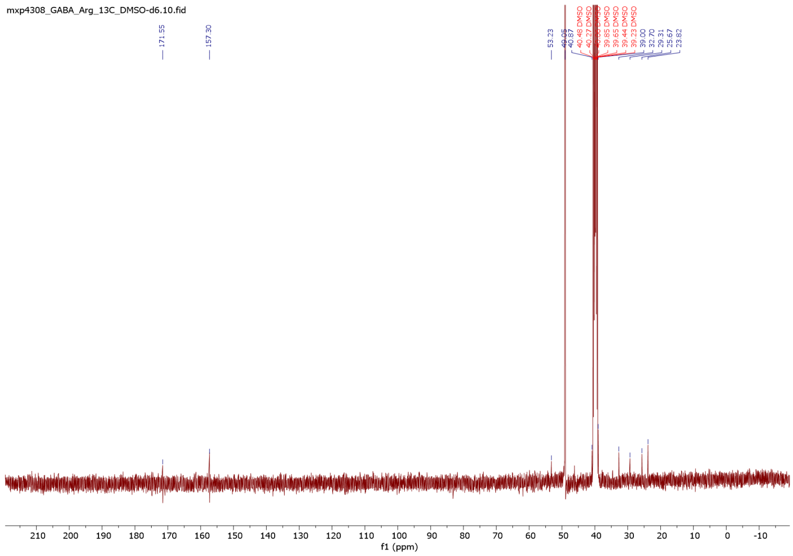

**Supplementary Fig. S6**: $^{13}C$ spectra of **2** (400 MHz, DMSO-d$_6$): $\delta$ 171.5, 157.3, 53.23, 39, 32.7, 29.31, 25.7, 23.82.

### Appendix S5 Elucidation of mental depression biomarkers

We used ICEBERG to hypothesize structure annotations for all MS/MS features with $p < 0.05$ from Nurses' Health Study and narrowed down to five proposed structures considering cost and human confidence. Two standards, hercynine (**1**) and LPC 19:0 (**9**), were confirmed by RT and MS/MS match, as reported in Fig. 3 and Extended Data Fig. 5. The following standards were ordered, but no RT match was observed:

1. For an unknown with RT = 5.94 min and precursor m/z = 146.08112, ICEBERG's top-5 prediction, 3-(Methylamino)-4-oxopentanoic acid (SMILES: CC(=O)C(CC(=O)O)NC), was ordered, but the RT of the standard did not match the unknown.
2. For an unknown with RT = 7.62 min and precursor m/z = 170.10034, ICEBERG's top-1 prediction, [(5-Methylthiophen-2-yl)methyl](propan-2-yl)amine (SMILES: CC1=CC=C(S1)CNC(C)C), was ordered, but the RT of the standard did not match the unknown.
3. For an unknown with RT = 8.19 min and precursor m/z = 130.08617, ICEBERG's top-2 prediction, 5-methylproline (SMILES: C[C@H]1CC[C@H](N1)C(=O)O), was ordered, but the RT of the standard did not match the unknown.

### Appendix S6 Elucidation of TB meningitis biomarkers and cost estimation of custom synthesis

The following compounds in Supplementary Table S17 were ordered for elucidating GABA-Arg (**2**) with an average cost of $2,100. When carrying out this structural elucidation campaign, in parallel with ICEBERG predictions, we used the popular structural elucidation tool, CSI:FingerID [36] in SIRIUS [37], to propose structures for the unknown with candidates from PubChem. The correct structure, GABA-Arg, was ranked at top 8 by SIRIUS. We also queried quotes for two possible dipeptides that consist of arginine and GABA, as ICEBERG's top-1 prediction suggests (Fig. 3e). The cost of custom synthesis in Fig. 1a is reported based on the statistics in Supplementary Table S17.

**Supplementary Table S17**: Custom synthesis for the elucidation of GABA-Arg.

| IUPAC name | Manufacturer | Amount | Cost |
|---|---|---|---|
| (2S)-2-(4-aminobutanoylamino)-5-(diamino methylideneamino)pentanoic acid* | AnaSpec Inc, Fremont CA | 10 mg | $564 |
| 4-[[(2S)-2-amino-5-(diaminomethylideneamino) pentanoyl]amino]butanoic acid | AnaSpec Inc, Fremont CA | 10 mg | $923 |
| (2S)-2-[[(2S)-2-aminopropanoyl]amino]-6-(diamino methylideneamino)hexanoic acid | ENAMINE US Inc, Monmouth Junction NJ | 100 mg | $3005 |
| (2S)-2-[[(2S)-2-aminopropanoyl]amino]-5-(diamino methylideneamino)pentanoic acid | ENAMINE US Inc, Monmouth Junction NJ | 100 mg | $3005 |
| (2S)-2-amino-5-[4-(diaminomethylideneamino) butanoylamino]pentanoic acid | ENAMINE US Inc, Monmouth Junction NJ | 100 mg | $3005 |

The verified structure, GABA-Arg (**2**), is the first entry marked by *. Any amount above 1 mg is sufficient for most MS/MS standards, while a custom synthesis company usually requires a minimum amount for placing an order. ICEBERG has the potential to save considerable budgetary costs, having ranked the correct structure at top-1. (Fig. 3e)

### Appendix S7 Pesticide degradation experimentation

#### S7.1 Experimental information

We investigated the abiotic breakdown of thiophanate methyl under controlled laboratory conditions. To do this, we prepared stock solutions of the pesticide at 10 $\mu$g/L by serially diluting a high-purity analytical standard with Milli-Q water. These solutions were distributed into six 30 mL quartz vials—three wrapped in aluminum foil to serve as dark replicates and three left unwrapped for light exposure. Two additional vials containing only Milli-Q water acted as blanks, one foil-wrapped and one unwrapped. All vials were placed in a solar simulator to replicate natural sunlight while maintaining uniform temperature and agitation, as adapted from [108]. Samples were withdrawn at 0, 0.5, 1, 2, 3, 7, 15, 25, 36, 50, and 70 days and analyzed by LC-MS/MS.

#### S7.2 LC-MS/MS procedure

For tandem mass spectrometry, analyses were performed using an Orbitrap mass spectrometer (Q Exactive, ThermoFisher Scientific), equipped with electrospray ionization (ESI) operated in positive mode. The spectrometer was calibrated using a standard calibration mix (Pierce™ FlexMix™ Calibration Solution from ThermoFisher Scientific) using a ThermoFisher procedure for a low mass scan range of 70-400 $m/z$. Samples were introduced via a Dionex Ultimate 3000 coupled UPLC system. Column used: ZORBAX RRHD Eclipse Plus C18 Column, 2.1 $\times$ 100 mm, 1.8 $\mu$m with ZORBAX RRHD Eclipse Plus UHPLC Guard pre-column and Viper Adaptor attachment. Mobile Phase A: 0.1% formic acid (v/v) in Optima LC/MS-grade water, Mobile Phase B: 0.1% formic acid in Optima LC/MS-grade MeCN. Flow rate: 0.300 mL/min. Injection volume: 20 $\mu$m. Column temperature: room temperature. The UHPLC program is a linear gradient from 5% to 100% B over 14 minutes, then a 1-minute hold at 100% B, followed by a 1-minute linear gradient back to 5% B, and a 3-minute equilibration hold. Parameters included a spray voltage of 3.50 kV, capillary temperature of 256 °C, and sheath, auxiliary, and sweep gas flow rates set at 48, 11, and 2, respectively. Tandem MS/MS spectra were acquired using HCD at stepped normalized collision energies of 30%, 40%, and 60%. Five MS/MS scans were taken in each cycle, with apex triggering set to 1-10 seconds and dynamic exclusion set at 5 seconds. The mass range for full-scan acquisition was set from 150.0 to 450.0 $m/z$ with a resolution setting of 35,000 for both MS and MS/MS scans. XIC peak areas were normalized to $^{13}C$-labeled atrazine and simazine internal standards. The detailed XIC peak areas for carbendazim at each time step for each experimental replicate are reported in Supplementary Table S18.

**Supplementary Table S18**: Extracted ion chromatogram peak area for 192.0764 $m/z$ (carbendazim, **14**) normalized by internal standard at different time steps (Fig. 3c). "Rep." means replicate number. "n.d." indicates not detected.

| Condition | Rep. | Days | | | | | | | | | |
|---|---|---|---|---|---|---|---|---|---|---|---|
| | | 0 | 0.5 | 1 | 2 | 3 | 7 | 15 | 36 | 50 | 70 |
| Light | 1 | 83800 | 430100 | 387500 | 214800 | 122600 | 19810 | n.d. | n.d. | n.d. | n.d. |
| | 2 | 44240 | 405100 | 430400 | 384200 | 354400 | 226600 | 123500 | 25190 | n.d. | n.d. |
| | 3 | 55990 | 380000 | 382000 | 347400 | 308600 | 180200 | 114400 | 10240 | n.d. | n.d. |
| Dark | 1 | 83800 | n.d. | n.d. | 116600 | 116400 | 265900 | 218600 | 417700 | 565200 | 542800 |
| | 2 | 44240 | n.d. | n.d. | 96430 | 126000 | 253900 | 251600 | 473400 | 574300 | 564500 |
| | 3 | 55990 | n.d. | n.d. | 195800 | 186900 | n.d. | 235900 | 474300 | 597800 | 498900 |

### Appendix S8 Pooled C-N coupling experimentation

#### S8.1 General Information

All reactions were run at a scale of 1 mL with the amine as the limiting reagent at 0.1 M. All reagents were prepared as stock solutions in DMSO such that a transfer of 0.1 mL would aliquot the substance needed to meet the specified reactant concentration at the target reaction volume. In the pooled experiment, 0.1 mL of each amine stock solution was pooled before 0.1 mL of resultant mixture was transferred to the reaction vessel. Order of addition: copper iodide, l-proline, potassium carbonate, halide(s), and finally amine(s).

#### S8.2 General Procedure

All chemical reactions were conducted in dried glassware and set up in a fume hood exposed to air. All solvents and reagents were purchased from Sigma Aldrich and were used as received. Glass 1-dram vials were used as reaction vessels, fitted with a screw-cap with a Teflon-coated silicone septa, and magnetic stir bars. Initial reaction analysis was typically performed by thin-layer chromatography on silica gel or using a Waters I-class ACQUITY UPLC-MS (Waters Corporation, Milford, MA, USA) equipped with in-line photodiode array detector (PDA) and QDa mass detector (ESI positive and negative ionization mode). 10 $\mu$L sample injections were taken from acetonitrile solutions of reaction mixtures or products ($\sim$1 mg/mL). Column used: Waters Cortecs UPLC C18+ column, 4.6 mm $\times$ 50 mm with Waters Cortecs UPLC C18+ VanGuard pre-column, Mobile Phase A: 0.1% formic acid in Optima LC/MS-grade water, Mobile Phase B: 0.1% formic acid in Optima LC/MS-grade MeCN. Flow rate: 1.44 mL/min. Column temperature: 45 $^{o}$C. A 6-minute method was used. Thin Layer Chromatography was performed on 25 $\mu$m TLC Silica gel 60 F254 glass plates purchased from Fisher Scientific. Visualization was performed using ultraviolet light (254 nm). Caffeine was used as the internal standard.

For tandem mass spectrometry, the same analytical method was used as Supplementary Section S7. Some detailed instrumental configurations are different. Mobile Phase A: 0.1% formic acid in Optima LC/MS-grade water, Mobile Phase B: 0.1% formic acid in Optima LC/MS-grade MeCN. Flow rate: 0.300 mL/min. Column temperature: 45 $^{o}$C. A 60-minute method was used. Parameters included a spray voltage of 4.20 kV, capillary temperature of 320 °C, and sheath, auxiliary, and sweep gas flow rates set at 50, 15, and 3, respectively. The mass range for full-scan acquisition was set from 150.0 to 350.0 $m/z$ with a resolution setting of 35,000.

#### S8.3 Singleton experiments

We describe the details of singleton experiments as follows. PROD/IS means the product XIC peak area normalized by the internal standard XIC peak area.

N-(3-phenylpropyl)aniline

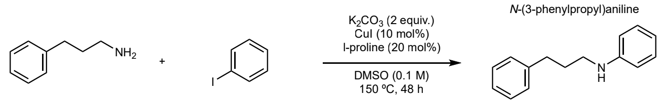

**17a** **18** (1.5 equiv.) **20a**

**Supplementary Fig. S7**: **17a** + **18** $\longrightarrow$ **20a**. To a 1-dram vial was added 0.01 mmol CuI (10 mol%), 0.02 mmol l-proline (20 mol%), 0.2 mmol potassium carbonate (2 equiv.), 0.15 mmol iodobenzene (**18**, 1.5 equiv.), and then 0.1 mmol 3-phenylpropan-1-amine (**17a**, 1 equiv.) before diluted to 1 mL with DMSO. The mixture was stirred on a hotplate at 150 ºC for 48 h before quenched, diluted with internal standard, and injected into an analytical column attached to a tandem mass spectrometer. The desired product (N-(3-phenylpropyl)aniline, **20a**) was found to elute at a retention time of 25.70 min. with a PROD/IS integration ratio of 8.27. HRMS (ESI): calculated $C_{15}H_{18}N$, $[M+H]^+$: 212.1434, found: 212.1431.

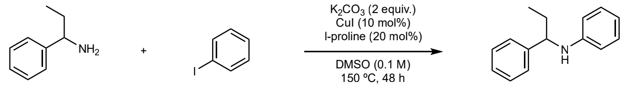

N-(1-phenylpropyl)aniline

**17b** **18** (1.5 equiv.) **20b**

**Supplementary Fig. S8**: **17b** + **18** $\longrightarrow$ **20b**. To a 1-dram vial was added 0.01 mmol CuI (10 mol%), 0.02 mmol l-proline (20 mol%), 0.2 mmol potassium carbonate (2 equiv.), 0.15 mmol iodobenzene (**18**, 1.5 equiv.), and then 0.1 mmol 1-phenylpropan-1-amine (**17b**, 1 equiv.) before diluted to 1 mL with DMSO. The mixture was stirred on a hotplate at 150 ºC for 48 h before quenched, diluted with internal standard, and injected into an analytical column attached to a tandem mass spectrometer. The desired product (N-(1-phenylpropyl)aniline, **20b**) was found to elute at a retention time of 34.82 min. with a PROD/IS integration ratio of 0.357. HRMS (ESI): calculated $C_{15}H_{18}N$, $[M+H]^+$: 212.1434, found: 212.1432.

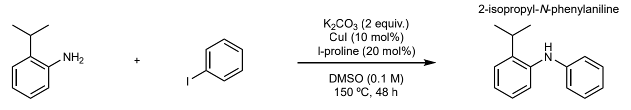

**17c** **18** (1.5 equiv.) **20c**

**Supplementary Fig. S9**: **17c** + **18** $\longrightarrow$ **20c**. To a 1-dram vial was added 0.01 mmol CuI (10 mol%), 0.02 mmol l-proline (20 mol%), 0.2 mmol potassium carbonate (2 equiv.), 0.15 mmol iodobenzene (**18**, 1.5 equiv.), and then 0.1 mmol 2-isopropylaniline (**17c**, 1 equiv.) before diluted to 1 mL with DMSO. The mixture was stirred on a hotplate at 150 $^{\text{o}}$C for 48 h before quenched, diluted with internal standard, and injected into an analytical column attached to a tandem mass spectrometer. The desired product (2-isopropyl-N-phenylaniline, **20c**) was found to elute at a retention time of 41.51 min. with a PROD/IS integration ratio of 2.48. HRMS (ESI): calculated $C_{15}H_{18}N$, $[M+H]^+$: 212.1434, found: 212.1431.

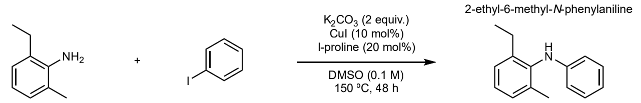

**17d** **18** (1.5 equiv.) **20d**

**Supplementary Fig. S10**: **17d** + **18** $\longrightarrow$ **20d**. To a 1-dram vial was added 0.01 mmol CuI (10 mol%), 0.02 mmol l-proline (20 mol%), 0.2 mmol potassium carbonate (2 equiv.), 0.15 mmol iodobenzene (**18**, 1.5 equiv.), and then 0.1 mmol 2-ethyl-6-methylaniline (**17d**, 1 equiv.) before diluted to 1 mL with DMSO. The mixture was stirred on a hotplate at 150 $^{\text{o}}$C for 48 h before quenched, diluted with internal standard, and injected into an analytical column attached to a tandem mass spectrometer. The desired product (2-ethyl-6-methyl-N-phenylaniline, **20d**) was found to elute at a retention time of 41.76 min. with a PROD/IS integration ratio of 0.683. HRMS (ESI): calculated C15H18N+ $[M+H]^+$: 212.1434, found: 212.1431.

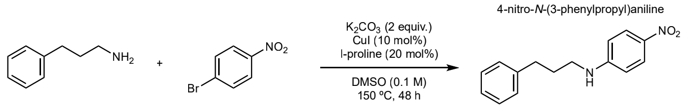

**17a** **19** (1.5 equiv.) **21a**

**Supplementary Fig. S11**: **17a** + **19** $\longrightarrow$ **21a**. To a 1-dram vial was added 0.01 mmol CuI (10 mol%), 0.02 mmol l-proline (20 mol%), 0.2 mmol potassium carbonate (2 equiv.), 0.15 mmol 1-bromo-4-nitrobenzene (**19**, 1.5 equiv.), and then 0.1 mmol 3-phenylpropan-1-amine (**17a**, 1 equiv.) before diluted to 1 mL with DMSO. The mixture was stirred on a hotplate at 150 $^{\text{o}}$C for 48 h before quenched, diluted with internal standard, and injected into an analytical column attached to a tandem mass spectrometer. The desired product (4-nitro-N-(3-phenylpropyl)aniline, **21a**) was found to elute at a retention time of 36.74 min. with a PROD/IS integration ratio of 3.00. HRMS (ESI): calculated $C_{15}H_{17}N_2O_2$, $[M+H]^+$: 257.1285, found: 257.1281.

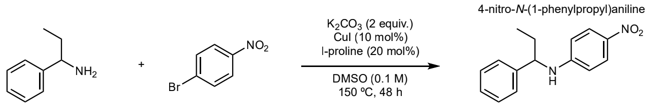

**17b** **19** (1.5 equiv.) **21b**

**Supplementary Fig. S12**: **17b** + **19** $\longrightarrow$ **21b**. To a 1-dram vial was added 0.01 mmol CuI (10 mol%), 0.02 mmol l-proline (20 mol%), 0.2 mmol potassium carbonate (2 equiv.), 0.15 mmol 1-bromo-4-nitrobenzene (**19**, 1.5 equiv.), and then 0.1 mmol 1-phenylpropan-1-amine (**17b**, 1 equiv.) before diluted to 1 mL with DMSO. The mixture was stirred on a hotplate at 150 ºC for 48 h before quenched, diluted with internal standard, and injected into an analytical column attached to a tandem mass spectrometer. The desired product 4-nitro-N-(1-phenylpropyl)aniline, **21b**) was found to elute at a retention time of 35.90 min. with a PROD/IS integration ratio of 0.147. HRMS (ESI): calculated $C_{15}H_{17}N_2O_2$, $[M+H]^+$: 257.1285, found: 257.1281.

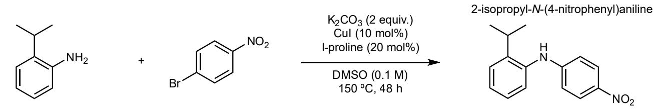

**17c** **19** (1.5 equiv.) **21c**

**Supplementary Fig. S13**: **17c** + **19** $\longrightarrow$ **21c**. To a 1-dram vial was added 0.01 mmol CuI (10 mol%), 0.02 mmol l-proline (20 mol%), 0.2 mmol potassium carbonate (2 equiv.), 0.15 mmol 1-bromo-4-nitrobenzene (**19**, 1.5 equiv.), and then 0.1 mmol 2-isopropylaniline (**17c**, 1 equiv.) before diluted to 1 mL with DMSO. The mixture was stirred on a hotplate at 150 ºC for 48 h before quenched, diluted with internal standard, and injected into an analytical column attached to a tandem mass spectrometer. The desired product (2-isopropyl-N-(4-nitrophenyl)aniline, **21c**) was found to elute at a retention time of 38.24 min. with a PROD/IS integration ratio of 0.454. HRMS (ESI): calculated $C_{15}H_{17}N_2O_2$, $[M+H]^+$: 257.1285, found: 257.1282.

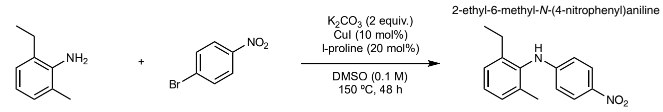

**17d** **19** (1.5 equiv.) **21d**

**Supplementary Fig. S14**: **17d** + **19** $\longrightarrow$ **21d**. To a 1-dram vial was added 0.01 mmol CuI (10 mol%), 0.02 mmol l-proline (20 mol%), 0.2 mmol potassium carbonate (2 equiv.), 0.15 mmol 1-bromo-4-nitrobenzene (**19**, 1.5 equiv.), and then 0.1 mmol 2-ethyl-6-methylaniline (**17d**, 1 equiv.) before diluted to 1 mL with DMSO. The mixture was stirred on a hotplate at 150 ºC for 48 h before quenched, diluted with internal standard, and injected into an analytical column attached to a tandem mass spectrometer. The desired product (2-ethyl-6-methyl-N-(4-nitrophenyl)aniline, **21d**) was found to elute at a retention time of 35.00 min. with a PROD/IS integration ratio of 0.213. HRMS (ESI): calculated $C_{15}H_{17}N_2O_2$, $[M+H]^+$: 257.1285, found: 257.1282.

#### S8.4 Pooled experiment

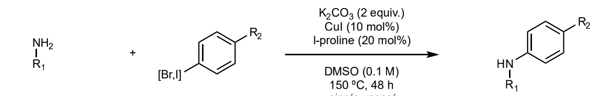

**17a-d** (total 1 equiv.) **18-19** (1.5 equiv.) **20a-d, 21a-d**

**Supplementary Fig. S15**: **17a-d** + **18** + **19** $\longrightarrow$ **20a-d** + **21a-d**. To a 1-dram vial was added 0.01 mmol CuI (10 mol%), 0.02 mmol l-proline (20 mol%), 0.2 mmol potassium carbonate (2 equiv.), 0.15 mmol iodobenzene (**18**, 1.5 equiv.), 0.15 mmol 1-bromo-4-nitrobenzene (**19**, 1.5 equiv.), and then 0.1 mmol amine pool (**17a-d**, total 1 equiv.) before diluted to 1 mL with DMSO. The mixture was stirred on a hotplate at 150 ºC for 48 h before quenched, diluted with internal standard, and injected into an analytical column attached to a tandem mass spectrometer. The desired products (**20a-d**, **21a-d**) were found to elute at a retention times of 25.70 min. (**20a**), 34.82 min. (**20b**), 41.51 min. (**20c**), 41.76 min. (**20d**), 36.74 min. (**21a**), 35.90 min. (**21b**), 38.24 min. (**21c**), and 35.00 min. (**21d**) with a PROD/IS integration ratios of 10.400 (**20a**), 0.322 (**20b**), 1.928 (**20c**), 0.450 (**20d**), 4.390 (**21a**), 0.328 (**21b**), 0.428 (**21c**), and 0.147 (**21d**). HRMS (ESI): calculated (**20a-d**) $C_{15}H_{18}N$, $[M+H]^+$: 212.1434, found: 212.1431 (**20a**), 212.1432 (**20b**), 212.1431 (**20c**), 212.1431 (**20d**); calculated (**21a-d**) $C_{15}H_{17}N_2O_2$, $[M+H]^+$: 257.1285, found: 257.1281 (**21a**), 257.1281 (**21b**), 257.1482 (**21c**), 257.1482 (**21d**).

### Appendix S9 Applying ICEBERG in novel 3-body reaction discovery

We add that the deconvolution of reaction products is not limited to instances of pooled reactions or crudes. Oftentimes understudied or novel reactions may generate a product with an ambiguous structure that can fit any number of structural hypotheses that match the identified mass. While this occurs less frequently in the case of bimolecular reactions (consider substrates with multiple reactive handles), multicomponent reactions can result in a diversity of potential product structures that are hard to predict due to their combinatorial nature. Fortunately, in these instances, isobaric product candidates are usually distinguishable by their fragmentation patterns. We exemplify this using a case study generated in a multicomponent reaction discovery campaign (Supplementary Fig. S16). In this reaction, a mass hit of interest was found during our rapid MS1 analysis, but isolation and 1-D NMR failed to confidently differentiate the true structure against a variety of possibilities, and ultimately, 2-D NMR was needed for full elucidation. However, ICEBERG was able to identify the true product structure using the fragmentation pattern generated using our MS/MS workflow, indicating its utility in a more rapid characterization technique in reaction discovery workflows. The experimental setup can be found in Mahjour et al. [33], with the 1-D and 2-D NMR spectra in the supplementary information (Figures S18-S21).

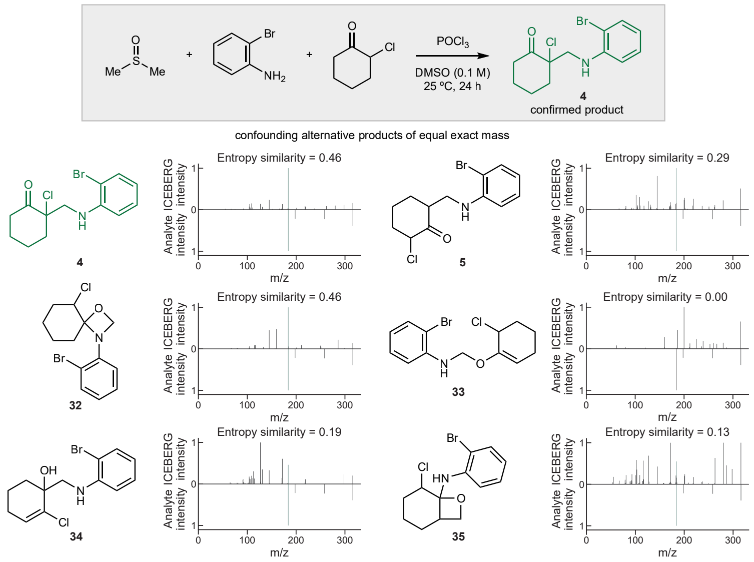

**Supplementary Fig. S16**: **ICEBERG deconvolutes isobaric potential products in the discovery of a novel 3-body reaction**. All six compounds shown (**4, 5, 32, 33, 34, 35**) are predicted as potential reaction products for the reaction shown in the gray box using the virtual flask developed by Mahjour et al. [33]. Among them, **5** cannot be distinguished from the confirmed product **4** by 1-D NMR, while ICEBERG successfully deconvolutes them by identifying the different fragmentation patterns of these two compounds. ICEBERG also predicts similar fragmentation patterns for **32** while 1-D NMR could tell it apart from **4**.

### Appendix S10 Enzymatic pathway modification site identification

#### S10.1 General information on elucidation of key intermediates

To elucidate the sequence of oxidative steps in the withanolide biosynthetic pathway, candidate genes were identified through *W. somnifera* genome sequencing and biosynthetic gene cluster analysis. Candidate cytochrome P450s (*CYP87G1*, *CYP88C7*, *CYP88C8*, *CYP749B2*) and the short-chain dehydrogenase *SDH2* were functionally characterized by stepwise heterologous expression in engineered yeast strains. Each strain expressed a defined combination of pathway enzymes to enable mapping of specific CYP functions:

- A strain expressing *CYP87G1* converted 24-methyldesmosterol to a C22-hydroxylated product (compound **24**).
- Co-expression of *CYP88C7* with *CYP87G1* yielded an additional C1-hydroxylated intermediate (compound **26**).
- Addition of *CYP749B2* resulted in successive C26 oxidations leading to lactone formation (compounds **27**–**28**).
- Inclusion of *SDH2* completed oxidation to the stable lactone (compound **29**).

Structures of key intermediates (**24**, **26**, **29**) were confirmed by NMR, as reported in ref. [91] (Supplementary Figures 6-11 for **24**, Supplementary Figures 13-18 for **26**, Supplementary Figures 19-24 for **29**). The sequential appearance of distinct oxidized intermediates upon stepwise enzyme addition established the functional order of *CYP87G1*, *CYP88C7*, and *CYP749B2* in the withanolide biosynthetic pathway.

For each modified strain, cultures were collected via centrifugation and resuspension in 500 $\mu$L from 3 mL samples of incubated media. Two series of extraction, involving the addition of 500 $\mu$L ethyl acetate, vortexed, and centrifuged at 21,130 rcf to separate layers and extraction of the organic layer (250 $\mu$L) were performed. Following a drying step and methanol reconstitution, 60 $\mu$L of supernatant was extracted for LC-MS/MS analysis. Tandem mass spectrometry analyses were performed using a high-resolution Orbitrap Exploris 120 mass spectrometer (Thermo Fisher Scientific) equipped with a heated electrospray ionization (H-ESI) source in positive ionization mode. Samples were introduced with a Vanquish Flex Binary UHPLC system equipped with a Vanquish Variable Wavelength Detector (Thermo Fisher Scientific). Solvent A was water with 0.1% formic acid and solvent B was acetonitrile with 0.1% formic acid. Column used: Kinetex C18 column, 2.6 $\mu$m particle size, 150 × 3 mm, 100Å (Phenomenex). Injection volume: 2 $\mu$L. Flow rate: 0.500 mL/min. Column temperature: 30°C. The following chromatoagraphy gradient was used: 0-2 min, 40% B; 2-7 min, 40-95% B; 7-12 min 95% B; 12-15 min, 40% B. The mass range for full-scan acquisition was set from 300.0 to 600.0 $m/z$ with data-dependent acquisition of the four highest intensity precursor ion, and was set with a resolution setting of 120,000, HCD collision energy of 30%, RF lens of 70%, and static spray voltage of 3500 V.

Further and supplementary details regarding the methods may be found in ref. [91].

#### S10.2 Elucidation of 25, a side product of 24 catalyzed by CYP88C8

In addition to the structures and intermediates identified as part of the main biosynthetic routes of **30** and **31**, a second product matching the mass of hydroxylated **24** was collected, noted as **25**. **25** is hypothesized to be a side product and was isolated from a yeast strain expressing *CYP87G1* and *CYP88C8*. The same modification site identification strategy with ICEBERG was applied to the spectra of **24** and **25**, to identify 6 possible hydroxylation sites on the A or B rings of the steroid nucleus. This structure, as a side product of the main biosynthetic pathway, was not reported in ref. [91]; as such, the exact modification was determined with $^1H$ NMR (collected from a 700MHz Bruker Avance Neo NMR spectrometer equipped with a Prodigy CryoProbe), which can be found in Supplementary Figures S17 and S18.

The NMR data for structural verification of **25**: $\delta$ 5.61 (1H, dd, $J$ = 1.9, 5.5 Hz, H-6), 3.86 (1H, m, H-7), 3.77 (1H, dt, $J$ = 2.5, 11.1, H-22), 3.59 (1H, m, H-3), 1.71 (3H, s, H-26), 1.71 (3H, s, H-27), 1.66 (3H, s, H-28), 1.00 (3H, s, H-19), 0.99 (3H, d, $J$ = 6.2, H-21), 0.73 (3H, s, H-18).

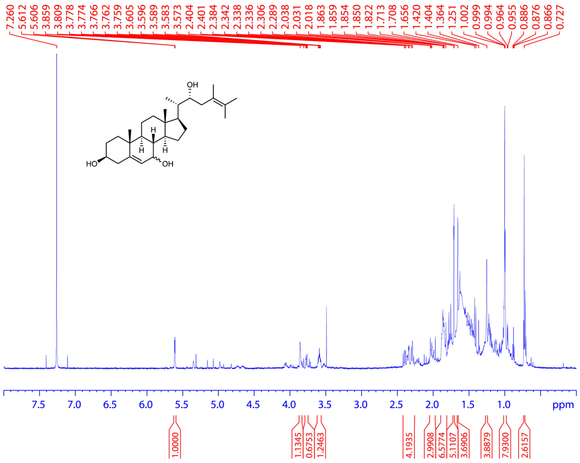

**Supplementary Fig. S17**: $^1H$ spectrum of **25**. (700 MHz, $CDCl_3$, 298 K).

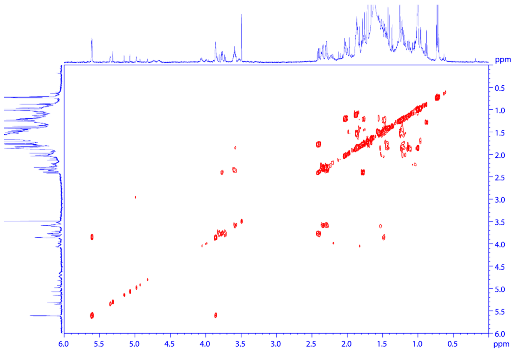

**Supplementary Fig. S18**: HHCOSY spectrum of **25**.
